# PLAE web app enables powerful searching and multiple visualizations across one million unified single-cell ocular transcriptomes

**DOI:** 10.1101/2022.09.12.507632

**Authors:** Vinay Swamy, Zachary Batz, David McGaughey

**Affiliations:** Department of Biomedical Informatics, Columbia University; Neurobiology, Neurodegeneration & Repair Laboratory, National Eye Institute, National Institutes of Health; Bioinformatics Group, Ophthalmic Genetics & Visual Function Branch, National Eye Institute, National Institutes of Health

**Author notes:** Correspondence: David McGaughey < >.

## Abstract

**PURPOSE:** To create a high performance reactive web application to query single cell gene expression data across cell type, species, study, and other factors.

**METHODS:** We updated the content and structure of the underlying data (single cell Eye in a Disk, scEiaD) and wrote the web application PLAE (https://plae.nei.nih.gov) to visualize and explore it.

**RESULTS:** The new portal provides quick visualization of over a million individual cells from vertebrate eye-and body transcriptomes encomopassing four species, 60 cell types, six ocular tissues, and 23 body tissues across 37 publications. To demonstrate the value of this unified pan-eye dataset, we replicate known neurogenic and cone macula markers as well as propose six new cone human region markers.

**CONCLUSION:** The PLAE web application provides the eye community a powerful and quick means to test hypotheses related to gene expression across a highly diverse, community-derived database.

## Introduction

The retina is composed of six major cell populations: the photoreceptors, horizontal, bipolar, amacrine, ganglion, and non-neuronal cells.^1^ These six populations can be further divided into dozens of cell types.^2^ Furthermore, human disease of the eye can derive from damage to specific cell types. Glaucoma is characterized by damage to the retinal ganglion cells while cone and cone-rod dystrophies are characterized by photoreceptor degeneration. Efforts to untangle the transcriptomics of these diverse cell types with single cell transcriptomics date to the early 2000s when researchers^3–7^ used RT-PCR approaches to amplify select transcripts in small numbers of cells. Later, researchers used microarray platforms to analyze a broader array of transcripts^8–14^

The modern era of single cell transcriptomics ha been marked by the introduction of droplet-based technologies, which enabled quantitation of thousands of cells in a single experiment. This approach was pioneered by Macoksko et al. and enabled the profiling of gene expression in 44,808 single cells in the adult mouse retina..^15, 16^ The commercialization of the droplet approach has enabled broad access to this technology and the ocular community have responded by publishing several dozen ocular studies using single cell transcriptome technology since Macosko publication.

These publications’s foci includes, but are not limited to, studying developmental transcriptomic dynamics, identifying genes to distinguish core cell types, proposing finer gradations of cell types within existing categories, and comparing ocular transcriptomes between organisms. Independent querying of these datasets requires a laborious series of steps.

Unfortunately manuscripts can only present a sliver of the underlying data and further exploration is laborious, technically challenging, and computationally expensive. Briefly, one would have to identify the relevant publication, find the associated data deposit, download the cell - count matrix, find the cell label table (and optionally the t-SNE or UMAP coordinates), and then load the data into R or python to run the desired queries. If one desired to compare across publications, then one would also have to align the cell type labels and should re-create the count quantification with consistent bioinformatic tooling. It is desirable, perhaps even necessary, to use multiple datasets to assess gene expression to confirm whether a gene expression pattern is consistent and reproducible. Reliance on a single dataset can be problematic as there can be technical issues - for example the large consortium dataset GTEx has persistent contamination from highly expressed genes across tissues.^17^

Web-based resources to query gene expression can democratize access to large datasets as the computationally intensive steps above can be run once and then shared to anyone with a internet connected device. Single cell RNA-seq web resources that contain ocular tissues include the UCSC Cell Browser, Tabula Sapiens, PanglaoDB, and the Protein Atlas.^18–21^ Primary limitations to these resources include incomplete cell type labels and sample metadata that make ocular focused queries challenging. In contrast to these, the Spectacle resource is an ocular specific web app which has curated over a million cells from dozens of independent resources.^22^ However each data deposit is independently processed, which makes cross study queries impossible. We recently created a unified ocular data resource which placed dozens of independent single-cell RNA-seq ocular and non-ocular datasets in a single combined space, the Single Cell Eye in a Disk (scEiaD).^23^

To broaden the usage of this powerful meta-atlas, we restructured the data into a custom sqlite-based database and built a reactive web application to access it. As many new studies were published since our last data build, we re-ran the scEiaD pipeline in March of 2022. In aggregate we have collated 44 datasets across 35 publications, four species, and 29 tissues. We hand-curated 60 cell type labels, totaling 606612 cells and used those ground-truth labels as the basis for a machine learning-based algorithm which labels the 500018 more cells. Quality control metrics were tuned and applied to remove lower quality cells. In the end, the single cell Eye in a Disk (scEiaD) database contains 1136041 cells and 1,509,179,347 non-zero gene/cell expression values. We demonstrate how this unified resource can be used to identify genes that consistently distinguish cell types across studies and species, find genes which have species-specific expression cell type patterning, demonstrate how our pre-computed differential testing can identify markers between similar progenitor cell types, and finally provide a R-based analysis document which end-to-end demonstrates how we use scEiaD to run a custom differential gene expression test to validate and propose new genes that distinguish macula from peripheral cones in humans.

## Methods

### scEiaD pipeline upgraded and the database re-built

The scEiaD pipeline is largely the same as in Swamy et al..^23^ Very briefly, we collected the raw fastq files and quantified with the kallisto / bustools system.^24, 25^ The quantified counts are merged into one matrix that is used for the downstream integration with scvitools (version 0.13.0) create a batch corrected lower dimensional space. The raw counts and scANVI batch corrected lower dimensional space is sent into Seurat for quality control, clustering, and 2D UMAP visualization. We use our previously published xgboost-based machine learning approach to transfer cell type labels to unlabelled cells.^23^

The following alterations were made: First, the cutoff to retain a cell was raised from 200 unique genes quantified to 300. Second, we use the DecontX algorithm^26^ from the celda R package (version 1.9.2) to automatically remove ambient RNA contamination on a per-study basis. Third, we altered the procedure for aligning gene names across species (see below for details). Fourth, we use the scANVI (scvitools based) batch correction method, which leverages the known cell type labels in the batch correction process.^27^

A custom cutoff for the minimum required number of unique genes identified per cell were used for the SRP362101 cornea dataset as an abnormally high number of cells were returned from our default value of 300. We set the cutoff for this studies to be 800 as this value resulted in approximately the same number of cells being returned as the authors reported in their papers.

We found that while nearly all datasets we curated could be integrated as a whole, a small number had strange behavior in the two dimensional UMAP space and/or nearly all cells from a study were machine labelled as a single cell type. These studies were hand-removed from the resource.

After evaluating the performance of the new dataset with our scPOP package we selected the following parameters for the integration with scANVI: 15 latent dimensions, 4000 highly variable genes, and 5 epochs. The UMAP was built with a min_dist of 0.1 and the clustering used 50 nearest neighbors.

### Cross species gene alignment

As we have four species in scEiaD (*Gallus gallus*, *Macaca fascicularis*, *Mus musculus*, and *Homo sapiens*) that we unify, we must identify shared genes. We found homologs/orthologs by pulling from the BioMart database “Ensembl Genes 105” and using *Homo sapiens* as the reference. We then selected orthologous gene names from *Gallus gallus*, *Macaca fascicularis*, and *Mus musculus*. This generates a table linked by Ensembl gene IDs and gene names. We then use a full join on non-duplicated Ensembl IDs between mouse and human. As we noticed a small number of gene names failed to be linked in this manner we ran another join on the remaining non-aligned genes, using gene name. This procedure gave us 17,769 shared human and mouse genes. To align the other species we again joined on Ensembl gene ID, removing genes where one chicken or macaque gene aligned to multiple human/mouse genes. In cases where multiple chicken or macaque genes aligned to one human/mouse gene, then we aggregated the counts by sum. In cases where there was no corresponding chicken or macaque gene, we filled in zeros. After these steps we had, as expected, 17,769 genes.

### Metadata curation

Every study brought into scEiaD was hand curated to identify unique biological samples, published paper ID (where available), organ, tissue (e.g. Cornea), source (iPSC based or tissue), single cell platform (e.g. 10Xv2, DropSeq, etc.). We also, where possible, annotated retina region (e.g. macula), sex, and age or developmental stage. Samples with perturbations (genetic or treatment) were not included. To import individual cell type labels (e.g. Rod, Muller Glia), we wrote custom code for each study that made this table available to link their cell type assignments back to the matching barcoded cell. We curated two types of cell type labels: “CellType” and “SubCellType” where the former is our normalized cell types taken from the published labels. We normalized names for CellType, for example changing “MG” to “Muller Glia” and also dropping more detailed cell type assignments like off and on bipolar cells for Bipolar Cells (as few studies went into this level). We did retain the original published (except for the broader name normalization) labels under SubCellType.

### Differential gene testing

We used a “pseudo-bulk” based approach where the counts for a category (CellType, Cluster, or CellType (Predict)) were summed for each category - organism - study grouping using the aggregateAcrossCells function from the scuttle R package.^28^ This created a counts matrix which has statistical properties approximating “bulk” RNA seq experiment matrices. This allowed the use of the more mature bulk RNA-seq tooling. This approach has also been found to be more specific in identifying differentially expressed genes in single cell RNA-seq experiments^29^. If there were fewer than 50 cells in organism - study accession - category combination, they were discarded for the differential test. Any genes with a sum of 0 counts after aggregation were removed. The aggregated matrix was imported into DESeq2 using the DESeqDataSetFromMatrix function with the design given as “∼study_accession + category.” Contrasts were extracted either with the target category (e.g. “Cone”) against all remaining or in a pair wise manner (“Cone” vs “Rod”) with the DESeq2 “results” function.^30^ We tested each species (Human, Macaque, Mouse) separately.

### PubMed Citation Search

The R package easyPubMed was used to search for 1000 randomly chosen genes present in scEiaD for both gene name alone and gene name plus “AND Retina” with the get_pubmed_ids function. We then used the returned Entrez identifier to pull the abstract with the fetch_pubmed_data function so we could extract the PubMed ID (PMID). This was repeated with the the genes identified as being well supported differentially expressed for ocular cell types. To assess whether the difference in distributions was statistically different, we used the base R t.test function.

### scEiaD structure and web app optimization

The web app at plae.nei.nih.gov can display gene expression across over a million cells in only a few seconds. This speed is only possible due to custom data structures that were optimized for data query efficiency. Like eyeIntegration’s EiaD database,^31^ scEiaD is a sqlite database with tables for the core categories. This allows the app to initialize in about 30 seconds on a cloud server with memory usage under 8 GB. For the gene by cell expression matrix, the data was transformed into a “long” format where there are three columns: gene, cell barcode, and gene count value. This allows for cell - gene level queries to be completed, again with minimal memory usage, in under 0.5 seconds. To generate the aggregated information (in the Dot Plot, Expression Plot, and tables) we pre-calculated all queries by running a dplyr “group by” operation on all column fields. On user given query, the data is further aggregated to the user request. This allows for complicated queries to complete in seconds rather than minutes. To compare web app loading times for useful tasks we queried https://plae.nei.nih.gov, http://singlecell-eye.org, and http://tabula-sapiens-portal.ds.czbiohub.org, on 2022-08-24. For Spectacle, we used the “Single-Cell RNA-Seq Analysis of Retinal Development…” 10X dataset from Clark et al. for all timings. Timings are accurate to the second. If the functionality tested was not available for a certain web app, it was represented by a blank space.

### Data reproducibility and availability

The code base for scEiaD is available at github.com/davemcg/scEiaD; the commit corresponding to this manuscript is #a99dced There are three relevant Snakemake pipelines which were used to create scEiaD: SnakeQUANT, which was used to quantify the gene / cell expression from the raw fastq files, SnakePOP, which runs the multi-parameter integration methods, and SnakeSCEIAD, which uses the optimal parameters identified in SnakePOP to run the cell type machine learning and differential gene testing that is incorporated into the scEiaD sqlite database for the app. The code base for the web app (along with local installation instructions) is available at github.com/davemcg/scEiaD/plaeApp. Data is available for download at plae.nei.nih.gov (click “Data”) and the Seurat object and metadata for the resource has been deposited at Zenodo with accession 10.5281/zenodo.7071682.

## Results

### New studies and improvements to the scEiaD database

Our first version of the single cell Eye is a Disk (scEiaD) database contained 34 studies, three species, 766615 cells, and 31 curated cell types.^23^ We updated the scEiaD database for the PLAE v0.91 web app in six important ways. First, we added a new species, chicken (*Gallus gallus*).^32^ Second we added two ocular outflow tract datasets, one brain choroid plexus, and three cornea datasets to enhance coverage across the eye.^33–38^ Third, we added a human pan-body reference scRNA dataset to allow for non-ocular comparisons.^39^ Fourth, we used an *in silico* method to remove background gene contamination as we noticed persistent Rhodopsin expression in many non-photoreceptors cells (Supplemental Figure 1). Fifth, we increased the cell retention cutoff for detected unique transcripts per cell from 200 to 300. Sixth, we carried over and harmonized several common cell type labels from the Tabula Muris project to more easily provide non eye cell type comparisons.^40^ As before, the pipeline’s scVI’s integration parameters were chosen rigorously by testing inegration performance across a wide range of latent dimensions and number of highly variable genes in our scPOP framework.^23^ Each study in the resource was hand assessed to determine whether they had any unusual properties in regards to machine learnt cell type proportions or position in the 2D UMAP space and a small number of outlier studies were removed from the resource.

After the dataset and quality control updates the scEiaD v2022-03-21 dataset now contains 44 studies, four species, 1136041 cells, and 60 curated cell types (Supplemental Figure 2). The scEiaD database contains 23 non-ocular human and mouse tissues and 6 ocular tissues across human, chicken, macaque, and mouse (Figure 1). This resource is only possible due to the publication of 35 single cell resources that scEiaD draws from (Supplemental Table 1). To recognize these papers, we prominently feature each of these studies along with a direct link to their citation, where possible, on the loading page of plae.nei.nih.gov.

**Figure 1:**
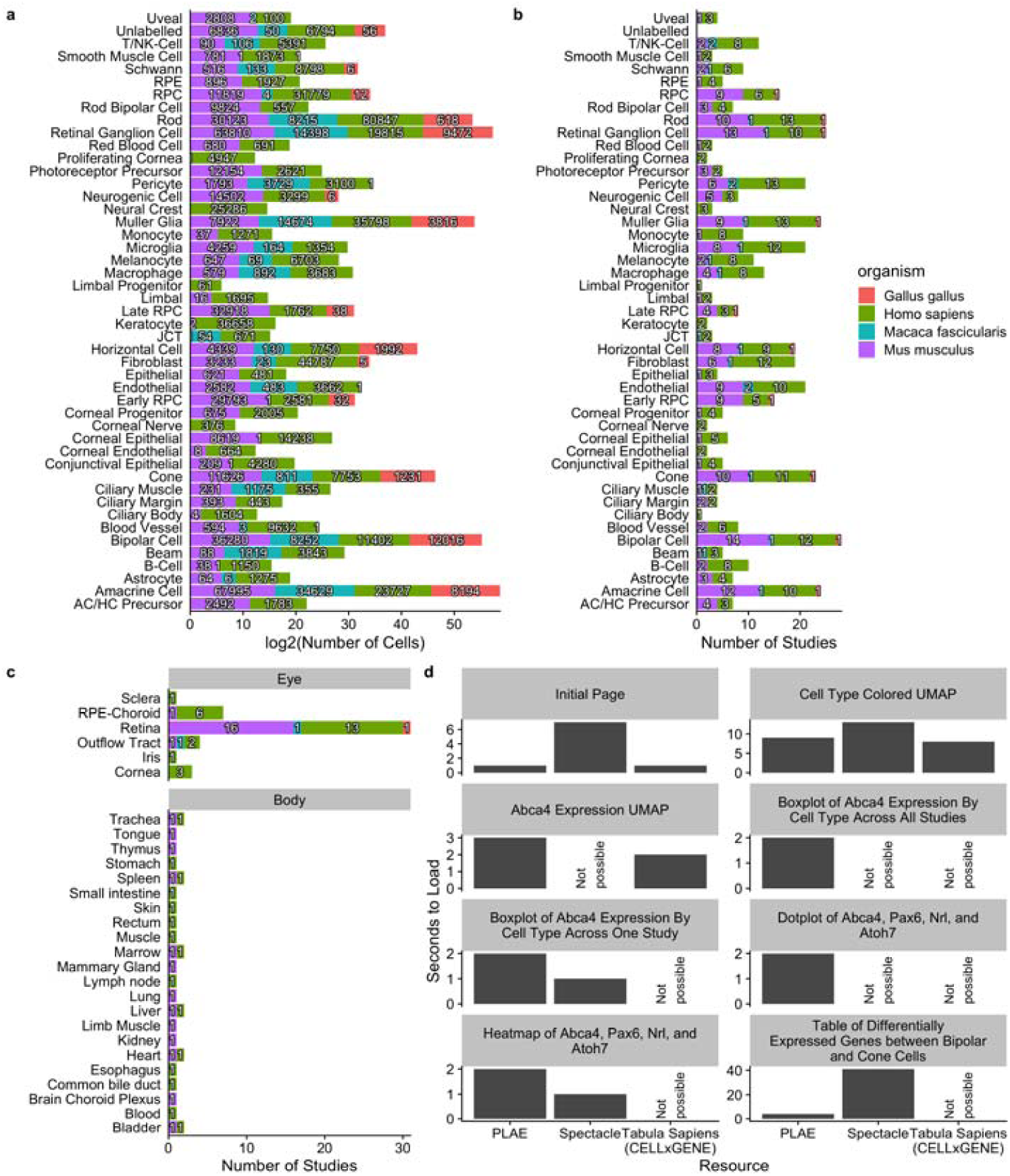
Distribution of tissues and cell types in scEiaD. A. Number of cells for each ocular cell type. B. Number of studies for each ocular predicted cell type (over 10 cells), colored by species. C. Tabulation of the number of studies present across 6 ocular and 23 non-ocular tissues, D. Web app timings (lower is better) for important functions compared between PLAE, Spectacle, and Tabula Sapiens (see methods for more details).

### PLAE resource contains several crucial features relative to other single cell resources

Several single cell RNA-seq web-based resources contain ocular data (Table 1). The UCSC Cell Browser, Tabula Sapiens portal, PanglaoDB, and Protein Atlas are not ocular focused, but do contain some ocular data. The PanglaoDB and UCSC are resources which have ingressed a large number of published single cell datasets The UCSC Cell Browser has around sixty thousand ocular single cell transcriptomes while the PangloaDB has about forty-seven thousand. However, these resources provide no curation of the metadata and process the data for each study individually. Thus each study is effectively silo-ed off from the rest and cross comparisons are impractical.

Table 1: Comparison of PLAE with other web resources that contain single cell ocular datasets, as of 2021-03-15. The Protein Atlas is asterisked as while it contains independent datasets, it only has one ocular study.

| Resource Name | Independent Datasets | Harmonization | Ocular and non-Ocular Data | Ocular Cell Count (thousands) | URL |
| --- | --- | --- | --- | --- | --- |
| PLAE | Yes | Yes | Yes | 1136 | https://plae.nei.nih.gov |
| Spectacle | Yes | No | No | 1500 | https://singlecell-eye.org |
| UCSC Cell Browser | Yes | No | Yes | 62 | https://cells.ucsc.edu |
| Tabula Sapiens | No | N/A | Yes | 10 | https://tabula-sapiens-portal.ds.czbiohub.org |
| PanglaoDB | Yes | No | Yes | 47 | https://panglaoDB.se |
| Protein Atlas | Yes* | No | Yes | 5 | https://www.proteinatlas.org/humanproteome/celltype |

The Tabula Sapiens project is an effort to sequence all cell type transcriptomes in humans; while it only contains around around ten thousand ocular cells it can be more readily cross compared to non-ocular tissues / cells. A more substantial downside is that the cell type labels can be insufficient - for example rods and cones are not independently identified but rather labelled as photoreceptors. The Protein Atlas resource has recently used only one retina single cell RNA-seq paper to provide cell type label focused information for gene queries (e.g. one can see whether a gene of interest is enriched in Cones). This is problematic as it is unclear whether signal found is specific to the one dataset they use or is common across independent species and datasets.

The Spectacle web resource is the closest comparison to PLAE, as it is a ocular focused resource which has collated a large number of ocular single cell RNA-seq resources (both organoid and tissue based datasets total well over a million cells) and made them available on a web app. PLAE is distinguished from these resources by hand curation of the cell and sample level metadata, re-processing of the data from the raw sequence in a consistent manner, integrating the independent studies to deliver cross study comparisons, and, crucially, providing full availability of the underlying data for outside usage.

While PLAE and Spectacle (https://singlecell-eye.org) both contain large numbers of ocular-related scRNA datasets, they are structured differently^22^. PLAE is built around the scEiaD database, which use a multi-stage pipeline to build and identify a high performing scVI-based single cell RNA eye model that is used to integrate all datasets together.^23^ This allows for gene queries to be constructed and analyzed across studies. In contrast, Spectacle, as of August 2022, is a compilation of study-level datasets and which makes queries across studies impossible. Spectacle does not contain any non-ocular datasets and the cell type labels have not been curated or harmonized between studies (for example different studies can use MG, Mueller, Müller Glia, or other variations to refer to the same cell type). Finally, in contrast to PLAE, Spectacle does not make their datasets available for download in any form.

### Extensive optimization makes the PLAE web app highly responsive despite containing huge amounts of data

There are two interwoven reasons why the PLAE app is quick, despite the huge amount of data it contains. First, the data structure of scEiaD is a sqlite database with pre-calculated tests stored in tables. A sqlite database and its indexing functionality allows large amounts of data to be stored for quick retrieval with minimal loading time and memory usage. The major downside is that this approach requires a large amount of disk storage, which can make local (on a personal computer) usage difficult; the scEiaD v2022-03-22 sqlite database is about 368 GB. Second, the visualization code in PLAE (e.g. the UMAP view) pull from the custom database directly and use a highly optimized plotting system (scattermore) to reduce drawing time. In contrast, starting from the loading page of Spectacle, it takes approximately eleven seconds to plot the cell type metadata for a several thousand cells across across a single study (tested on 2022-08-26). Within the same time, PLAE can plot the cell type labels for over a million cells across 34 studies (Supplemental Figure 2d). The speed of the PLAE app is also comparable to the CELLxGENE browser serving the Tabula Sapiens resource while providing additional functionality.

### Rich web-available visualizations

PLAE has several ways to visualize the data: one being the 2D UMAP view which, side-by-side, shows gene - cell expression and metadata - cell values. So thus, a user can see *Crx* expression across over a million cells and separately see cell type labels for those same million cells. The 2D UMAP view also has a zoom function, so a user can focus on a subset of cells. Clicking on the plot will show metadata for the five nearest cells to the click, which is useful to seeing what kind of cells are present in a small area (Figure 2a).

**Figure 2:**
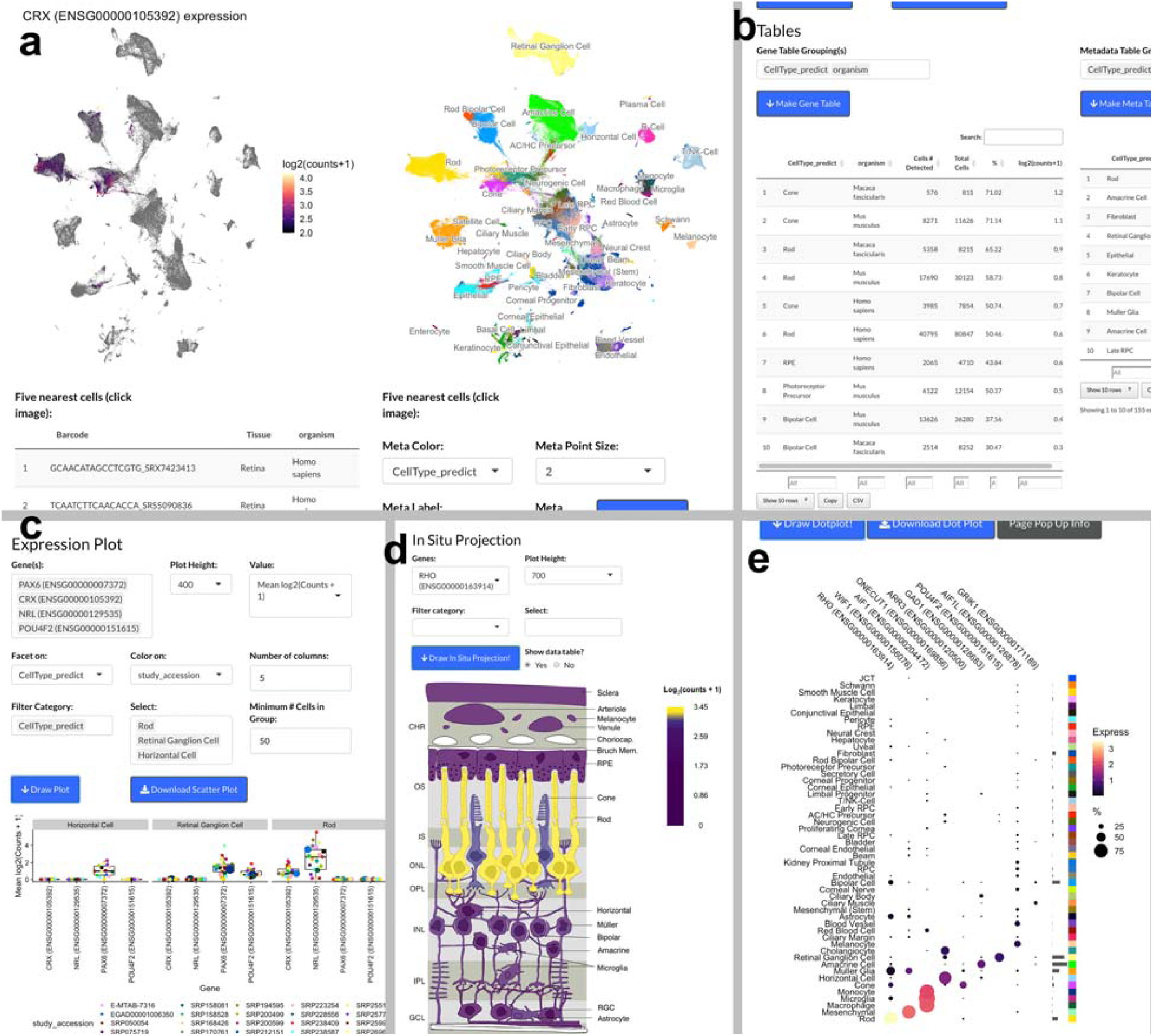
Screenshots from PLAE app. A. UMAP view showing Crx expression with cell type labels, B. Information table showing expression of Crx across celltypes and organism. C. Box plot of selected genes in a user selected cell types across different studies. D. In silico in-situ of Rho. E. Dotplot demonstating efficient display of gene expression across many cell types.

For users interested in seeing how expression of a gene relates to study or other covariates, the Expression Plot view displays expression by cell type, predicted cell type, or cluster. The values can then be split and colored by multiple covariates like study or organism. This allows a user to see whether a gene pattern is consistent across studies or species. We also offer a Dot Plot view which can display a large number of genes in a space efficient manner. This allows users to see whether a set of genes has related (or not) expression patterns across multiple cell types (Figure 2b and c).

While scEiaD does not have spatial scRNA seq datasets as of 2022, we do leverage existing knowledge of the structure of the retina to show *in silico* visualization of genes by cell type / layer in a cartoon cross section of the retina. The structure of the cartoon retina was based on the human retina. This unique visualization displays the major cell types of the retina and the RPE / choroid layer behind it. The cell types are colored by the relative amounts of expression in the user selected gene. Like all visualizations in PLAE, the user can use powerful filtering options to only display data from certain metadata characteristics, like species or publication (Figure 2d).

We have also added a Heatmap visualization which summaries the relative expression differences between user selected cell types or clusters across human, macaque, and mouse. This plot differs from the Dot Plot view as the expression values are relative instead of absolute. That is to say that while the Dot Plot view is proportionate to counts, the Heatmap view is showing the difference in expression between a given cell type (or cluster) and all other cells. This view is potentially useful if a user is especially interested in gene expression patterns across a variety of cell types.

### Well supported ocular cell types identify high confidence cell type markers

Cell type marker genes can be proposed on the basis of existing knowledge about their function and on high expression differences when comparing to other cell types. We can further narrow down the list of markers by using the high diversity of studies and organisms in scEiaD to propose a set of community supported cell type markers. We first identify a set of well supported cell types which we define as cell types that are detected in two or more independent studies across both human and mouse (Supplemental Table 3). These cell types were then assessed to identify differentially expressed genes which met the following criteria across human, mouse, and macaque: 1. padj less than 1×10^-4^ in two or more species, 2. mean log2 fold change greater than 2, and 3. mean padj less than 1×10^-5^. A final filter was applied to remove genes as candidates if they were differentially expressed (using the criteria above) in more than three different cell types. This left us with 2790 gene - cell type markers, a substantial reduction from the 33927 we had initially. We provide a table of these genes as supplementary file “consistently_differentially_expressed_genes.csv.gz.”

To visualize these top markers we select up to eight genes (ordered by the mean log2 fold change across organisms) per well supported cell type for each and plot the log2 fold change of the gene expression relative to all other cell types (Figure 3). We see that human, mouse, and macaque have similar expression patterning of the proposed markers across the well supported cell types. The cell types are arranged by the columns by how related the expression patterns are. We see how the differentiated cell types of the retina are grouped together with the exception of the Müller glia, which are near the RPC from which they are derived.

**Figure 3:**
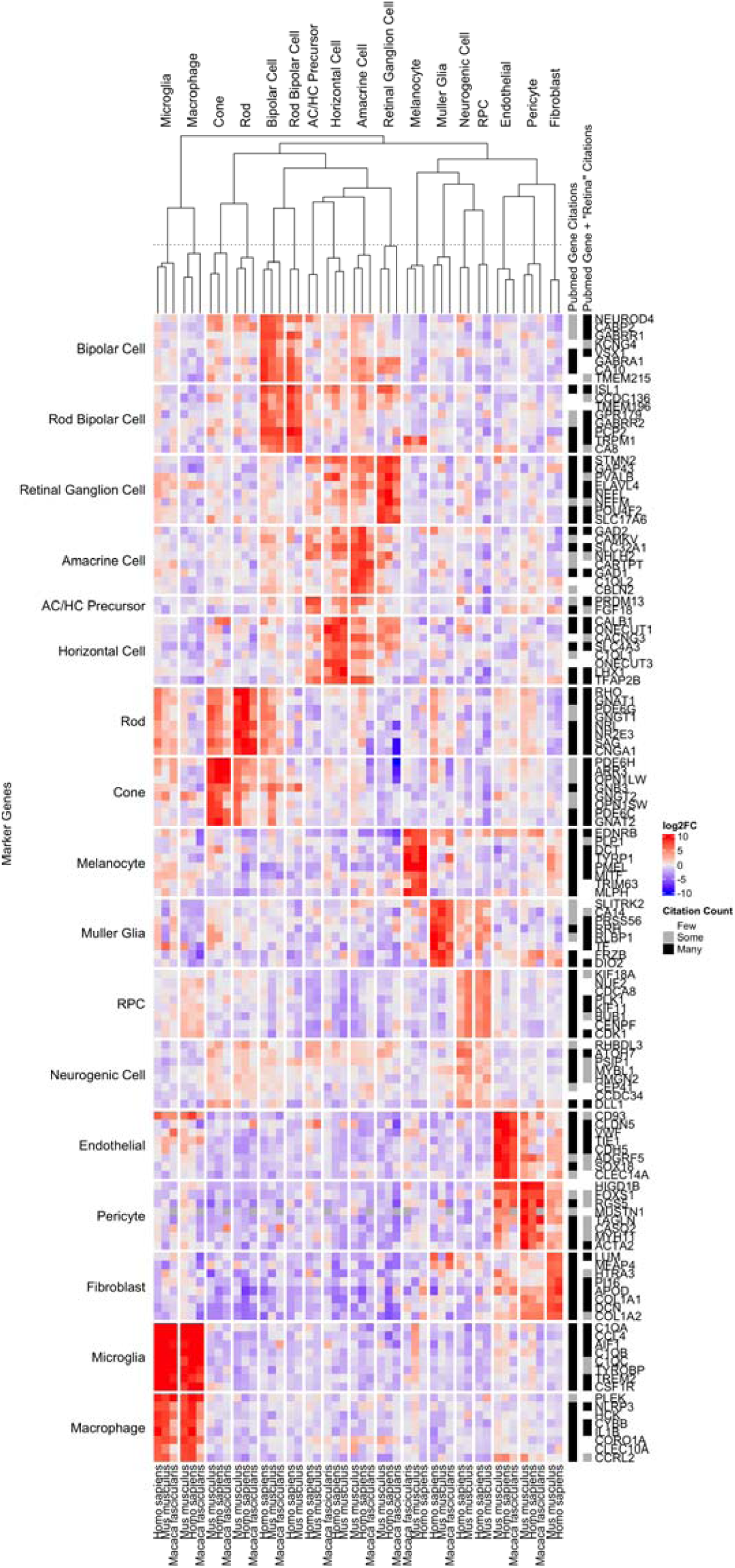
Rows are genes, columns are cell types and organisms. Rows are split by what cell type the gene is a marker for. Columns are split by the cell types. More intense red is a higher log2 fold change of the gene - cell type relative to all other cell types and blue is more negative. Relative number of citations are shown for the gene or gene ” and retina” when searching against PubMed compared to 1000 randomly chosen genes.

To assess whether this marker selection was identifying known or potentially novel marker genes we ran several sets of queries against PubMed. We searched for the gene name or the gene name plus ” AND Retina” across all well supported genes against the PubMed database, counting how many citations were returned. In contrast we also selected 1000 random genes that were not considered differentially expressed and ran the same PubMed searches. We find a substantial enrichment of citations for our well supported differentially expressed genes (t test, p < 6.7×10-98 and p < 2.4×10-12) for the gene and gene ” and retina” PubMed searches when compared against the 1000 random genes, respectively. To display the results of this search for each individual gene we show visually whether the number of citations returned is less or equal than the median citation count for the random gene set (“few”), more than the median (“some”), or more than the mean (“many”). We see that nearly all genes in (Figure 3) reflect existing knowledge.

Genes with few citations can be considered as novel markers. We see, for example, *Onecut3* appears to well separate the horizontal cells and has few PubMed citations. To assess whether there are any other good candidates with few citations, we produced another heatmap (Supplemental Figure 3). Most of these genes are differentially expressed across more than one cell type. We hand selected *C1ql2* and *Cartpt* (amacrine), *Frzb* and *Slitrk2* (Muller glia), and *Onecut3* (horizontal) and plotted these five genes in the boxplot view to confirm specificity across the full scEiaD database (Supplemental Figure 4). All five look to be specific to a cell type, except for *Frzb* which also is expressed in the RPE.

Other strategies for ranking genes are possible. We show in Supplemental Figure 5b and c how one can directly use our powerful web app to select candidate bipolar cell differentially expressed genes from the differential testing table and then quickly display the expression pattern of those genes in a compact heatmap visualization. The wealth of visualization options in plae allow for quick prototyping and hypothesis testing of gene expression patterning across many cells and studies.

### Tease apart the similar neurogenic and RPC cell types

While the well supported markers we propose clearly define most of the cell types, we see that the neurogenic cells and RPCs have very similar expression patterns across the gene sets. These cell types are closely related as the neurogenic cells are derived from the multi-potent RPC to later specify neural cell types. We use the set of genes identified by Lu et al.^41^ in their study of human retinal differentiation to determine whether our scEiaD database retains the same trends (Figure 4A). As expected, we see several patterns. First, we see most of the gene expression changes between RPC and neurogenic are matched between human. We further show that these genes also show the same trends in mouse datasets. Next we see that the direction of the changes largely matches Lu et al (Supplemental Figure 6). Finally, we see a small number of genes are confirmed in our database as consistently differentially expressed with a padj < 0.1.

**Figure 4:**
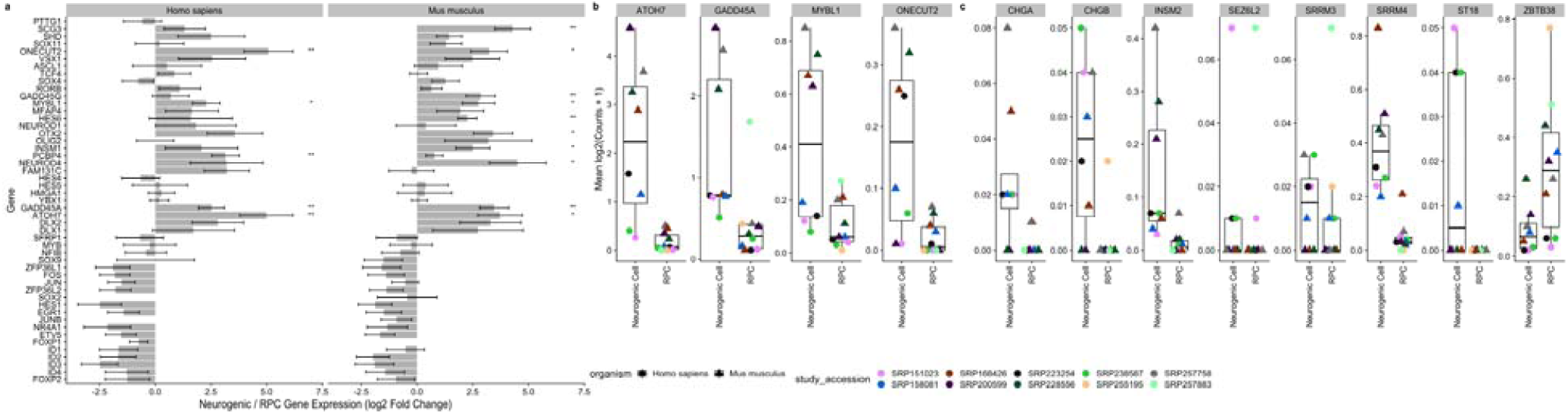
A. Genes proposed by Lu et al. as being significantly differentially expressed during the RPC to neurogenic differentiation in human. We plot the differential expression (log2(fold change) directly between RPC and neurogenic cells. The standard error (calculated by DESeq2) is plotted. Genes with a padj < 0.1 and padj < 0.01 are labelled with “*” and “**“, respectively. B. Genes from Lu et al. that we replicated with the scEiaD data. C. Novel genes we propose as being differentially expressed between RPC and neurogenic cells

To determine whether we can propose any new genes that distinguish RPC and neurogenic states we use our DESeq2 differential expression contrast that directly compares these two cell types. We find 88 human genes and 91 mouse genes that have an absolute log2 fold change greater than two and a adjusted p-value below 0.05 (Supplemental Table 4). To identify candidate genes that are consistently differentially expressed at this transitional state in mouse and human we apply similar logic that we used in identifying well supported marker genes across the retina cell types. We filter to genes with a padj less than 0.05 in both human and mouse, a mean log2 fold change greater than two or less than negative one (Figure 4B and C). We find one gene, *Zbtb38*, that meets this criteria and drops in expression when comparing RPC to neurogenic. We find 11 genes that increase in expression from RPC to neurogenic, 7 of which are not previously identified by Lu et al (Figure 4C). We replicate *Atoh7, Onecut2, Gadd45a, Mybl1* from Lu et al (Figure 4B) as differentially expressed between neurogenic and RPC. Futhermore we show in Supplemental Figure 5a how this figure can be replicated directly in our web app.

### Our pan-study macula vs peripheral cone test replicates a published macaque finding and proposes six new markers

**Figure 5:**
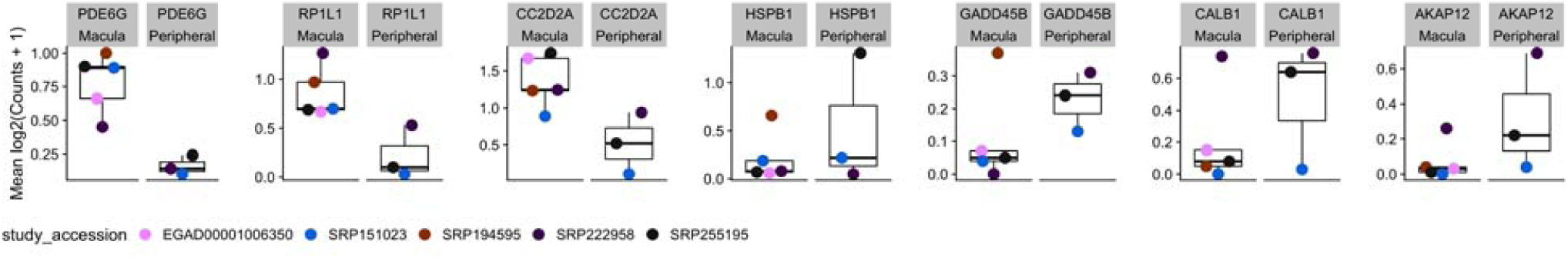
Plot of markers distinguishing human macula or peripheral punch derived cones.

While we have provided data in many forms via the web app at plae.nei.nih.gov, there are many more dissections of the data that can be performed. To facilitate custom testing with the scEiaD database we provide, as an example, a human-specific macula versus peripheral cone differential testing analysis document as supplementary file “pseudobulk_cone_region.freeze01.html.” In our analysis document we use the Seurat object containing the full scEiaD dataset we provide at plae.nei.nih.gov to build a custom pseudobulk counts matrix containing only cones from human studies that were taken in a region specific manner. This left us with 26678 cells across six studies. Yan et al. previously proposed *Prph2* and *Rs1* as markers to distinguish macula and peripheral located human cones^2^ and Peng et al. proposed *Calb1* and *Gngt1* as cone region markers in macaque.^42^

Our differential testing found that while *Prph2*, *Rs1*, and *Gngt1* had expression differences matching that the findings Yan et al. and Peng et al. published, they failed to replicate in our test with pvalue and padj being above 0.1 for all three. In contrast *Calb1* did replicate strongly with the sixth lowest padj (0.00721) across our entire test and a log2 fold change of 2.87. We find six more genes with a padj < 0.01: *Pde6g*, *Akap12*, *Hspb1*, *Gadd45b*, *Cc2d2a*, and *RP1l1* (Figure 6 and Supplemental Table 5). None of these candidate cone region marker genes display regionally different expression in rods (Supplemental Figure 7).

### Limitations of scEiaD

The longer term goal is to continue adding datasets to the scEiaD at plae.nei.nih.gov until we have three or more independent datasets per species and most ocular cell types. Earlier we proposed well supported cell types. Right now, most of these well supported cell types are in the retina. The RPE behind the retina, the cornea and lens in front, and the outflow tract and supporting musculature still could use more independent studies. Many common human diseases underlie the RPE (e.g. AMD) and the cornea (e.g. Keratoconus) and these tissues will remain a focus for the next update of our database. While we have curated a large number of cell types across the eye, many of these cell types have “sub” cell types. For examplethe human has three types of cones that are optimized for short, medium, and long wavelengths. More dramatically, Yan et al. have proposed over sixty different mouse amacrine cell types.^43^ While we have found it fairly straightforward to use machine learning to transfer the broad (e.g. rod, cone, amacrine) cell type labels across all datasets, we have found it very challenging to transfer the “sub” cell type labels. This is largely because while we have high diversity of studies for the broad cell types, this is not true for most of the sub celltypes. We believe more datasets that carefully dissect the sub-celltypes may be necessary to confidently propose the sub-celltypes of the retina. Finally, while our scVI based model can theoretically be used by an outside group to transfer their internal data into the scEiaD latent space, it is very challenging to implement as the the software landscape is still rapidly changing and it is very difficult to architect consistent outputs over time with regularly changing software versions. We are investigating simpler methods to enable transfer of knowledge from scEiaD to outside data.

## Conclusions

### High feature web app provides quick access to over one million transcriptomes across 44 datasets

We have improved on the scEiaD dataset we published in Swamy et al.^23^ by adding another species (chicken), new ocular tissues, a human non-ocular reference set; in total 10 new single cell datasets. We re-structured the data into a custom database to enable our custom built high performance web app to quickly access the data. This democratizes access to over a million cells across 44 studies to anyone with a web browser. We recognize that while we have built a powerful visualization platform we cannot anticipate all uses and for those researchers who wish to make custom queries of the data, we provide the data underlying PLAE in a wide variety of formats, including anndata and Seurat objects for each study, count tables, cell level metadata, and the DESeq2 based differential testing across three species. We provide a short example of how a novel query could work by running a custom differential test between macula and peripheral derived human cones and demonstrate how this can both validate existing proposed genes and suggest new ones.

### Diversity of studies enables reproduction and extension of cell type gene expression findings

A great many groups have proposed cell atlases. These range from efforts focusing on a subset of cells from a single layer of a tissue (for example^44^) to a huge consortium encompassing multiple organs.^18^ We propose a complementary approach wherein we use the research data from the community at large to create a meta-atlas. This approach enables a small group to produce an atlas which rivals the size from massive multi-group consortiums. Furthermore the inherent variability between the data from independent groups can be used as a feature to get insight into whether a gene-cell type signal is stable across an entire community’s datasets. We briefly demonstrate how the high study diversity in scEiaD can be used to validate proposed RPC to neurogenic gene expression changes.

## Supporting information

consistently_differentially_expressed_diff_table

consistently_differentially_expressed_genes

differential_expression_table

pseudobulk_cone_region_analysis

pseudoBulk_cone_region_table

R scripts

## Supplementary Figures and Tables

**Supplemental Figure 1:**
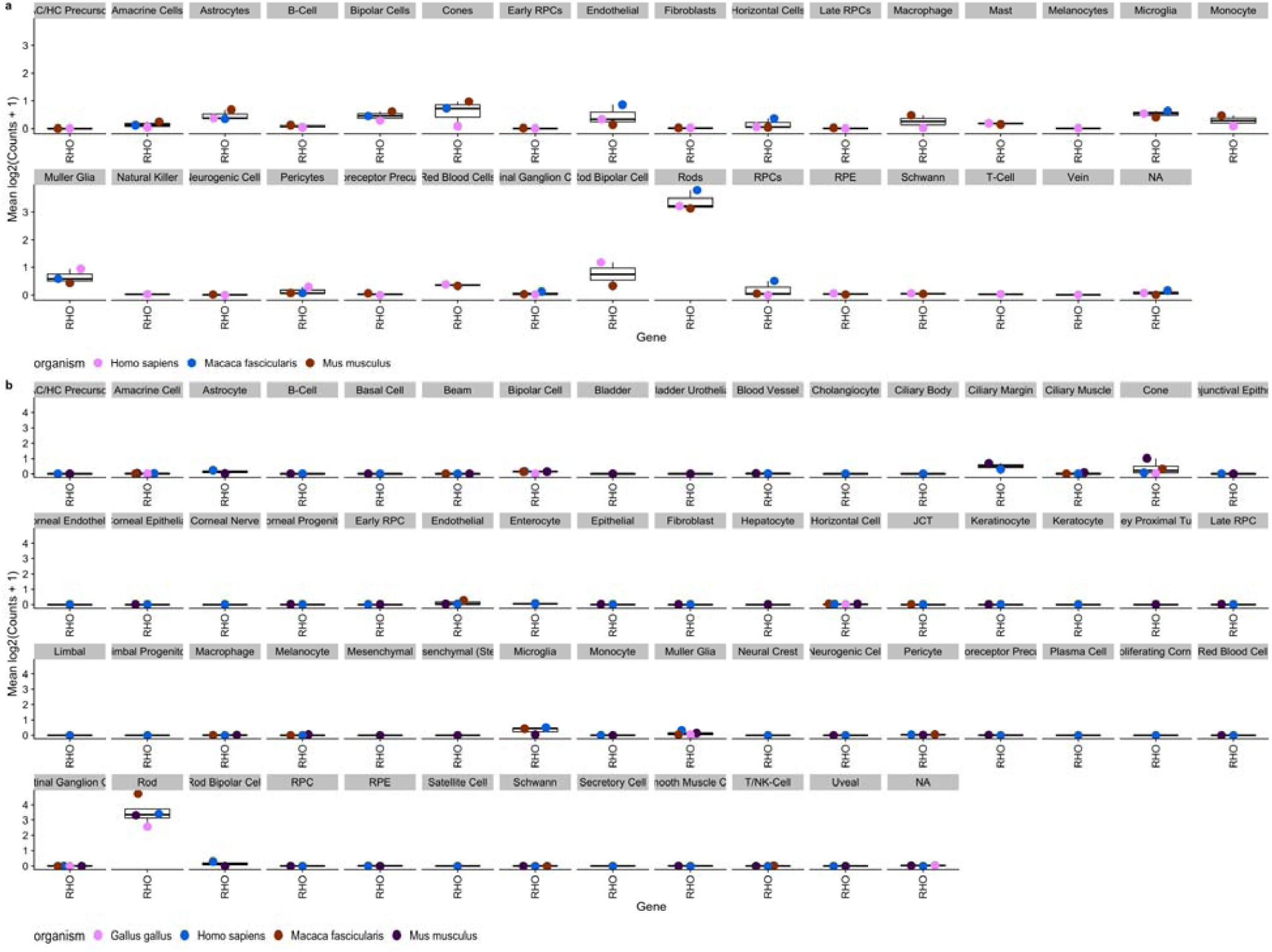
DecontX in silico ambient RNA contamination tool substantially removes Rhodopsin expression in non-rod cells while retaining high expression rods. First shown (a) is Rhodopsin expression across the scEiaD v0 dataset, without DecontX optimization. Note Rhodopsin expression is noticeable in many cell types beyond the rods. Next (b) is the scEiaD v1 dataset with DecontX optimization. Rhodopsin is nearly exclusively expressed in labelled rod cells. Despite the computational ambient RNA removal, Rhodopsin expression remains high in the Rods.

**Supplemental Figure 2:**
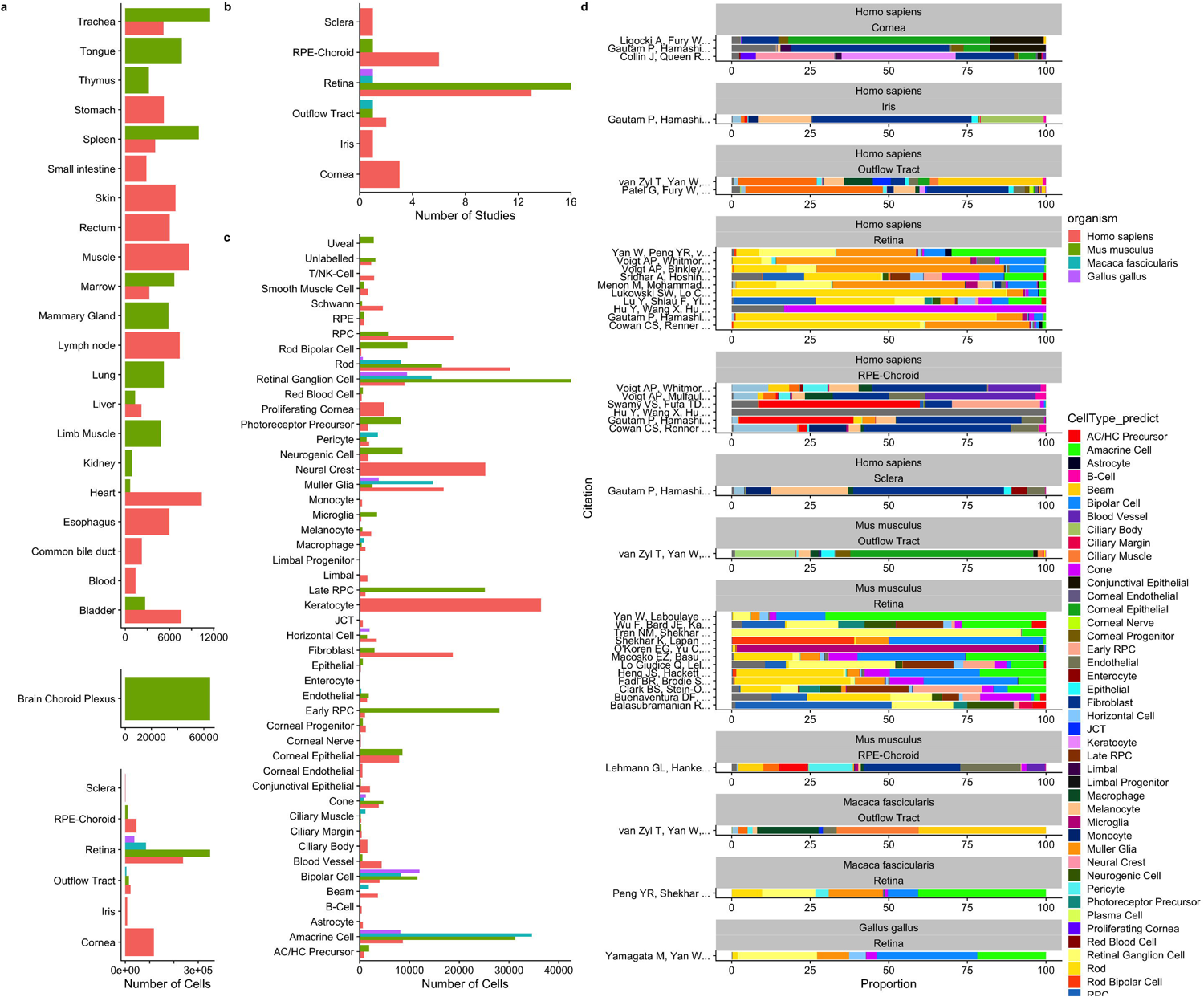
Distribution of tissues and cell types in scEiaD. A. tabulation of the number of cells present across 6 ocular and 23 non-ocular tissues, B. Number of studies for each ocular tissue. C. Number of cells present across 53 curated ocular-derived cell types. D. Proportion of cell types (predicted) across each study (split by species and tissue).

**Supplemental Figure 3:**
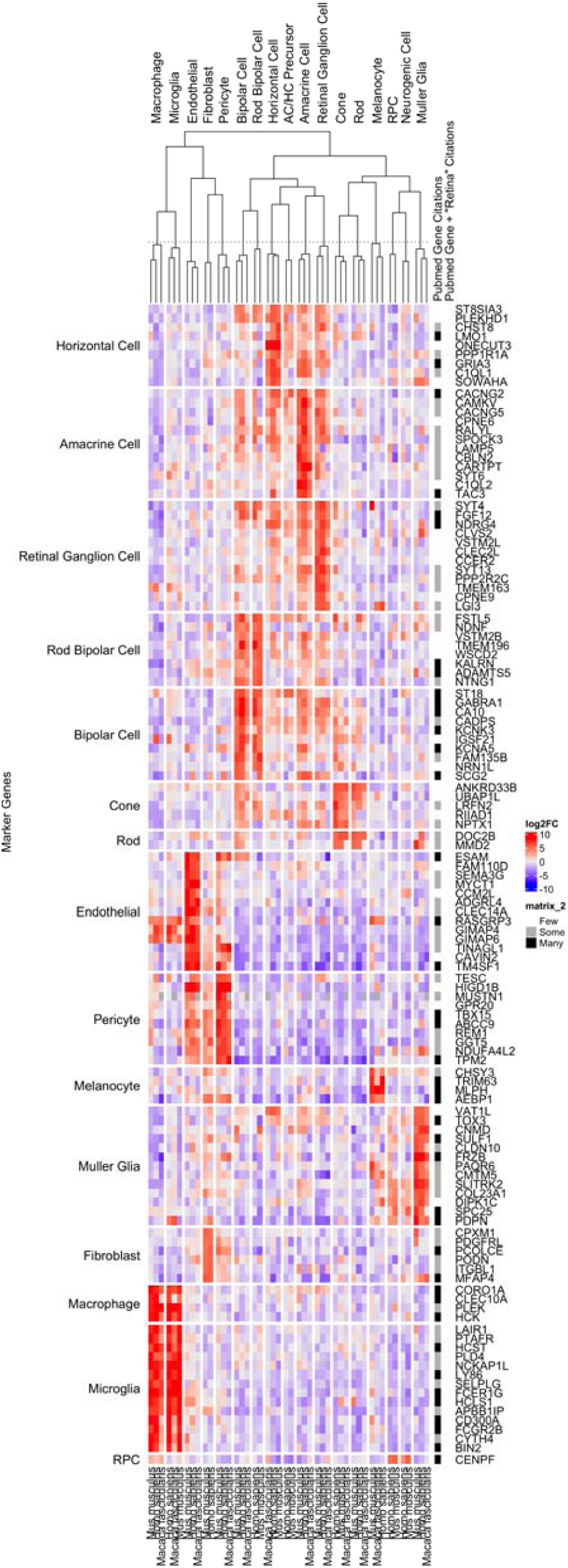
Well supported genes with a mean log2FC > 5 and zero pubmed citation hits for [gene] AND Retina search.

**Supplemental Figure 4:**
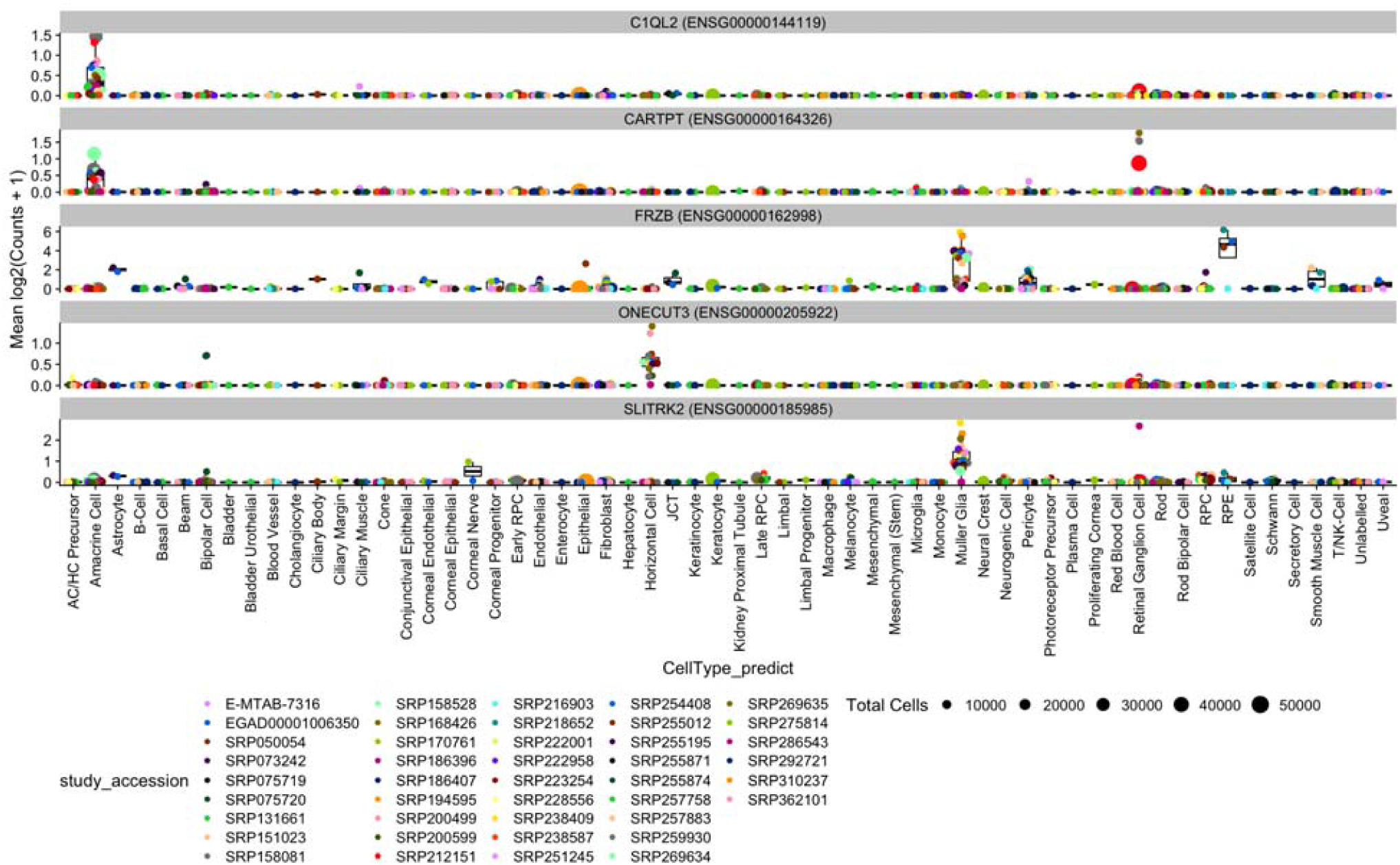
Five hand-selected genes selected from Supplementary Figure 3 as having fairly unique expression to a celltype while having few pubmed citations. Plotted as expression/boxplots to confirm specificity at the study level.

**Supplemental Figure 5:**
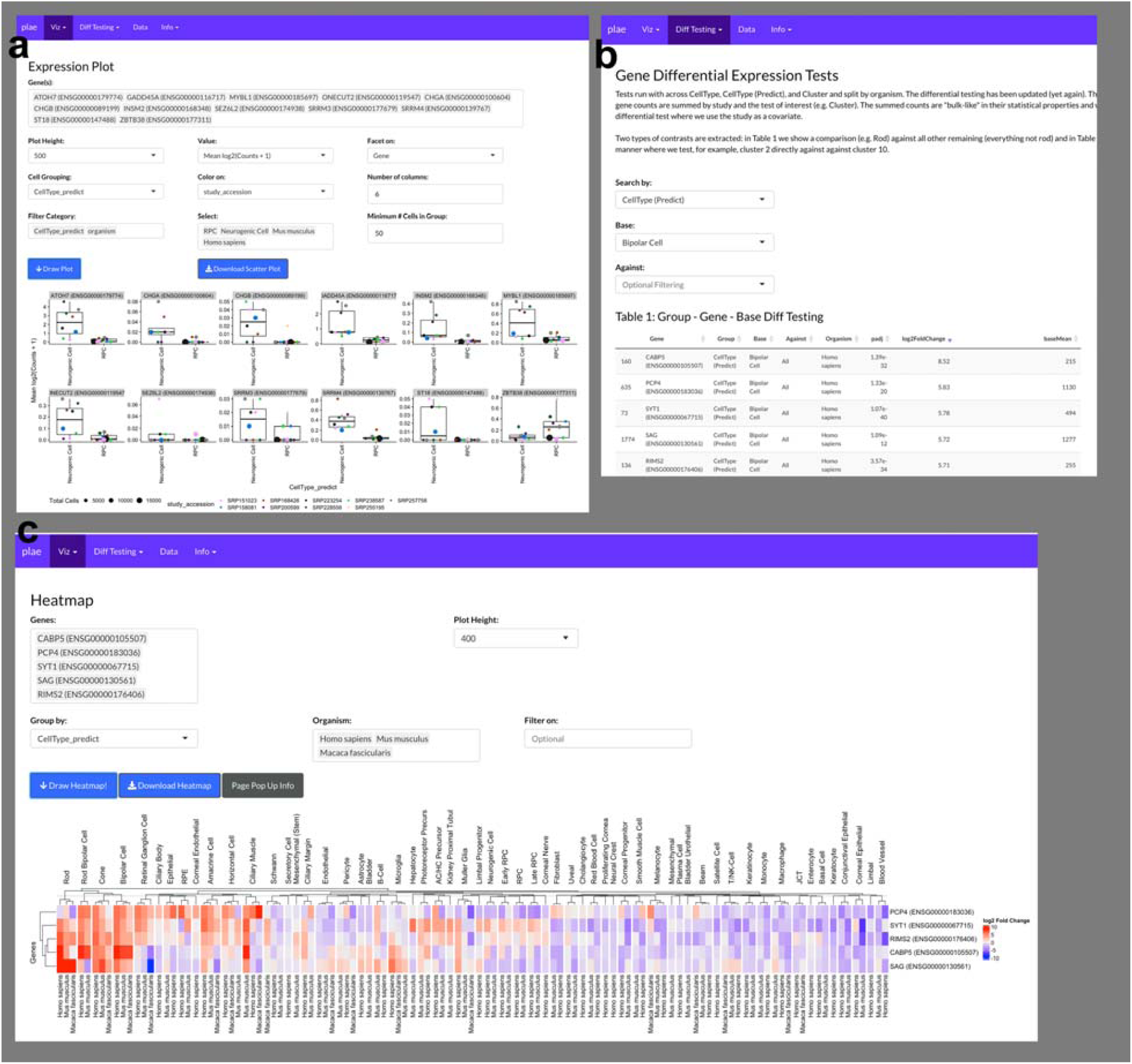
a. Recreation of Figure 4 directly in the plae.nei.nih.gov web app. b. Identify potential bipolar cell markers with the precomputed differential testing and in c. visualize the top five genes with a Heatmap.

**Supplemental Figure 6:**
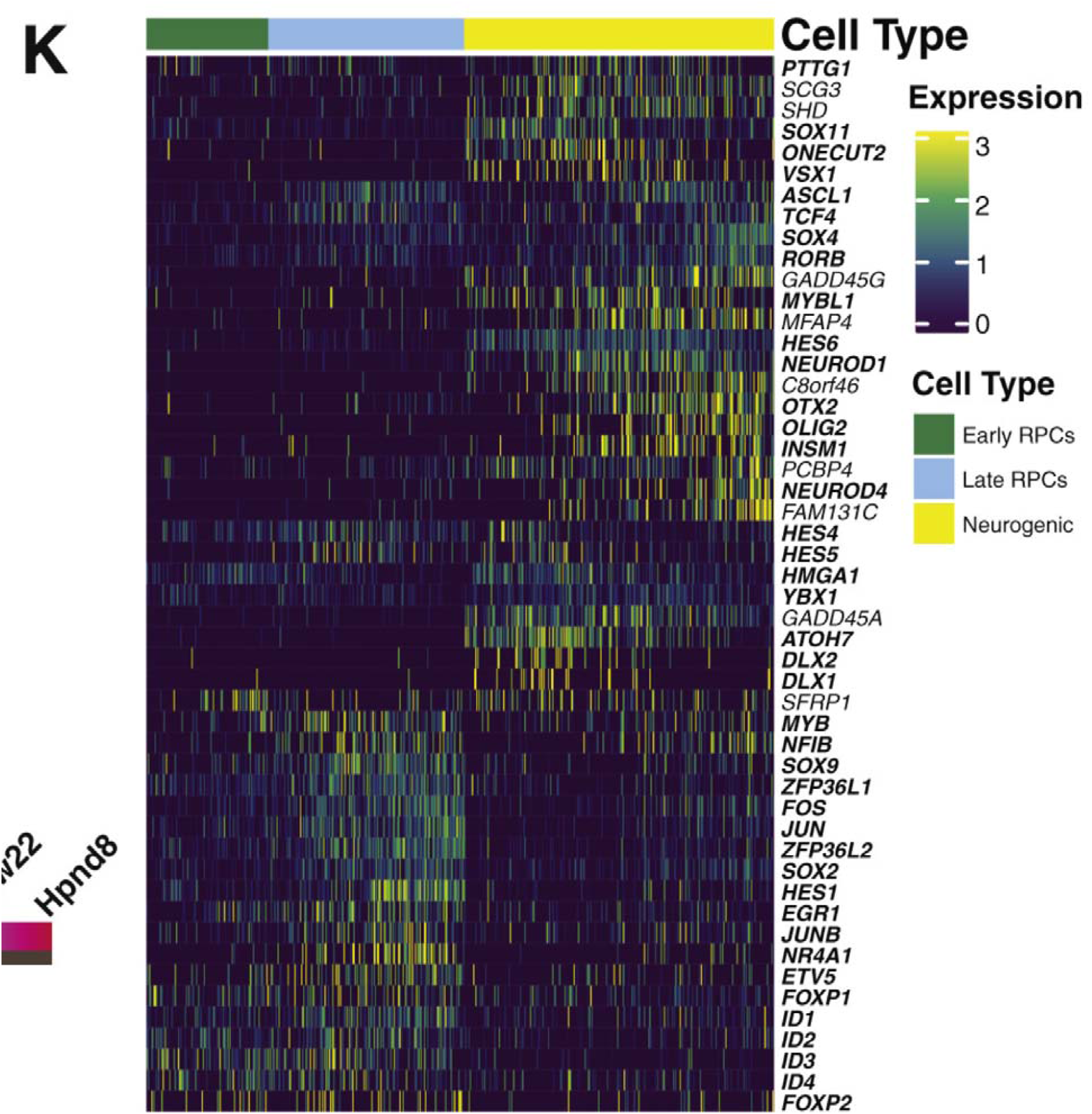
Supplemental figure 2K from Lu et al showing their candidate RPC / Neurogenic differentially expressed genes

**Supplemental Figure 7:**
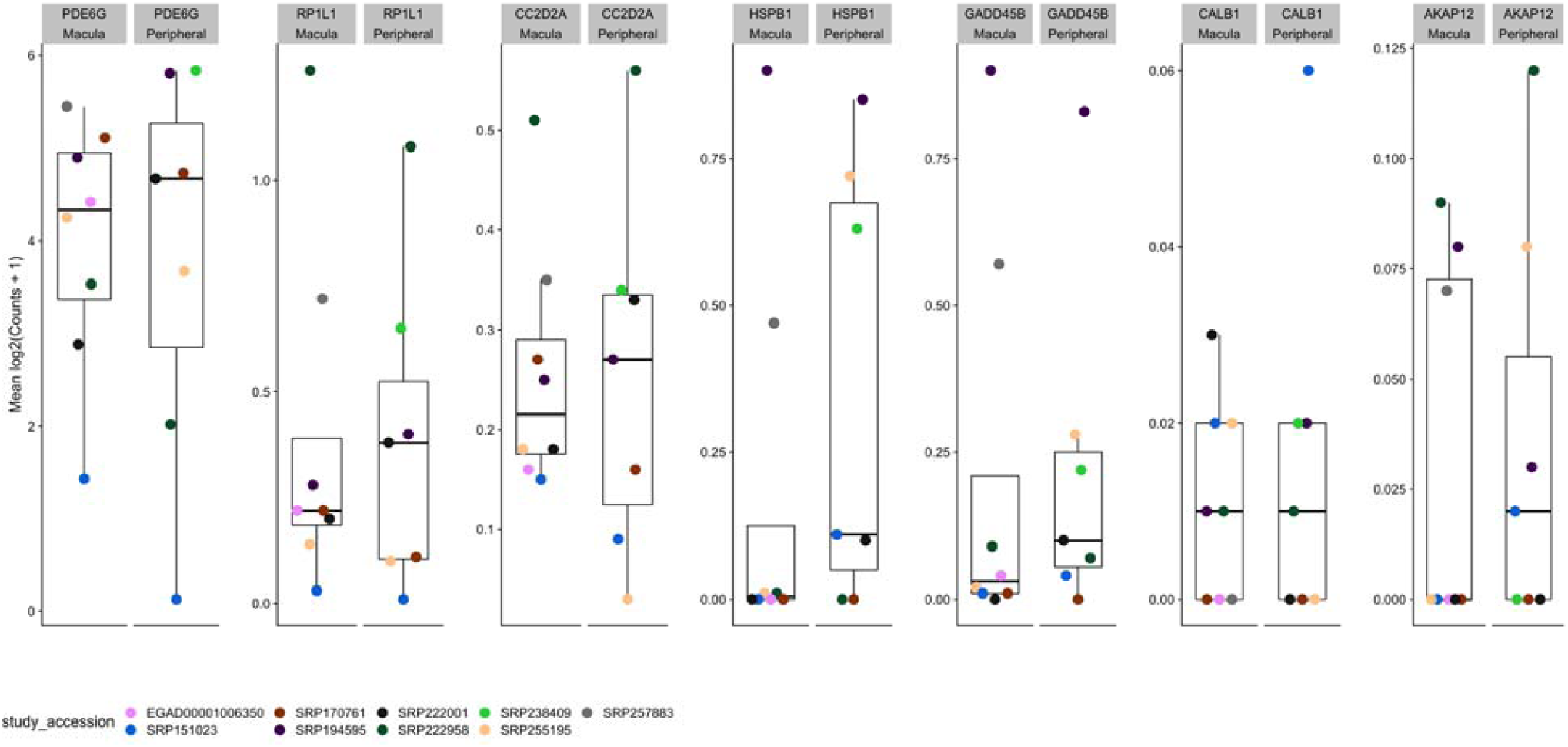
Human cone fovea - peripheral differentially expressed genes are not differentially expressed in human rods.

**Supplemental Figure 8:**
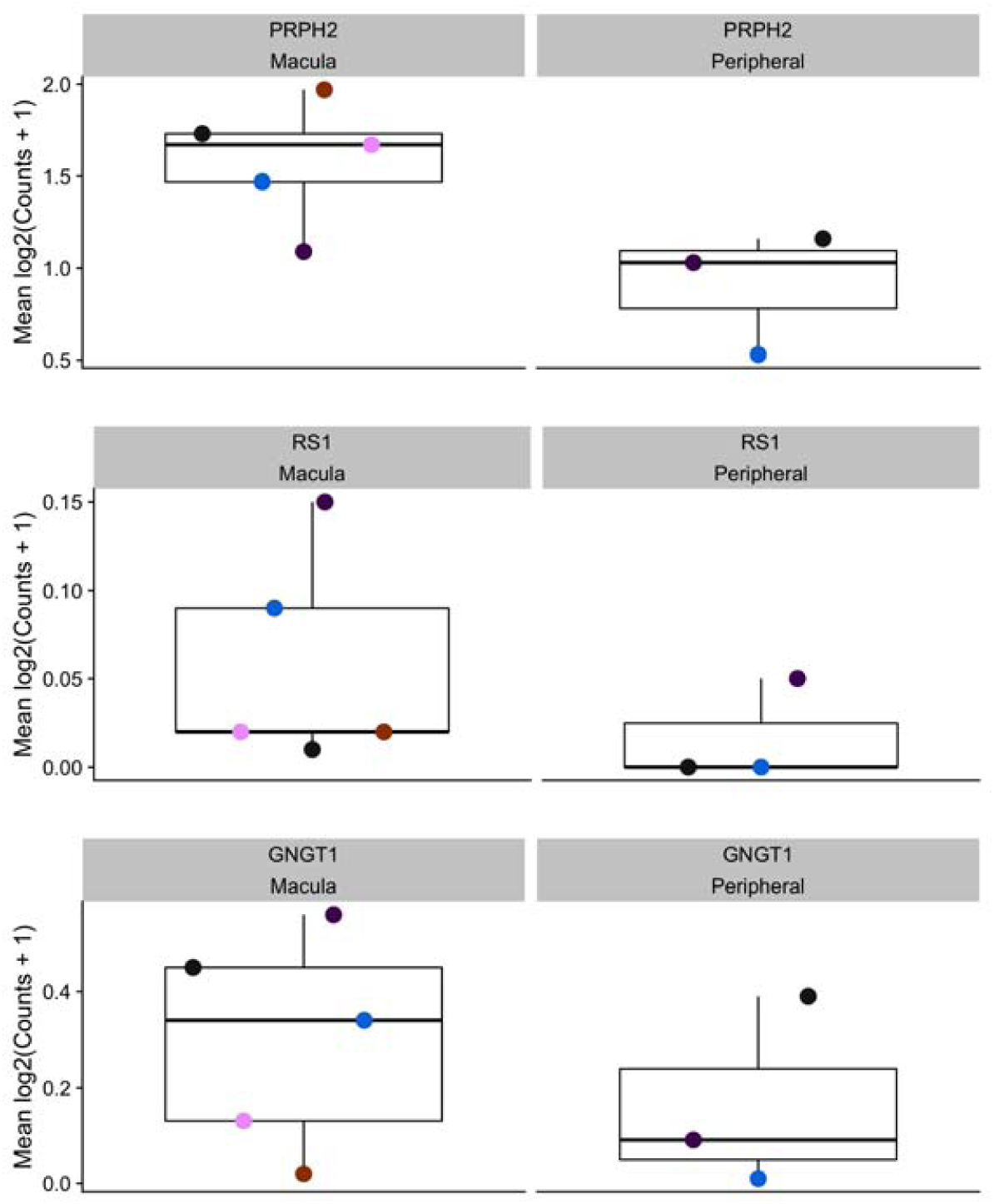
Genes identified by Peng et al. and Yan et al. as macula / peripheral differentially expressed in cones.

**Supplemental Table 1:**
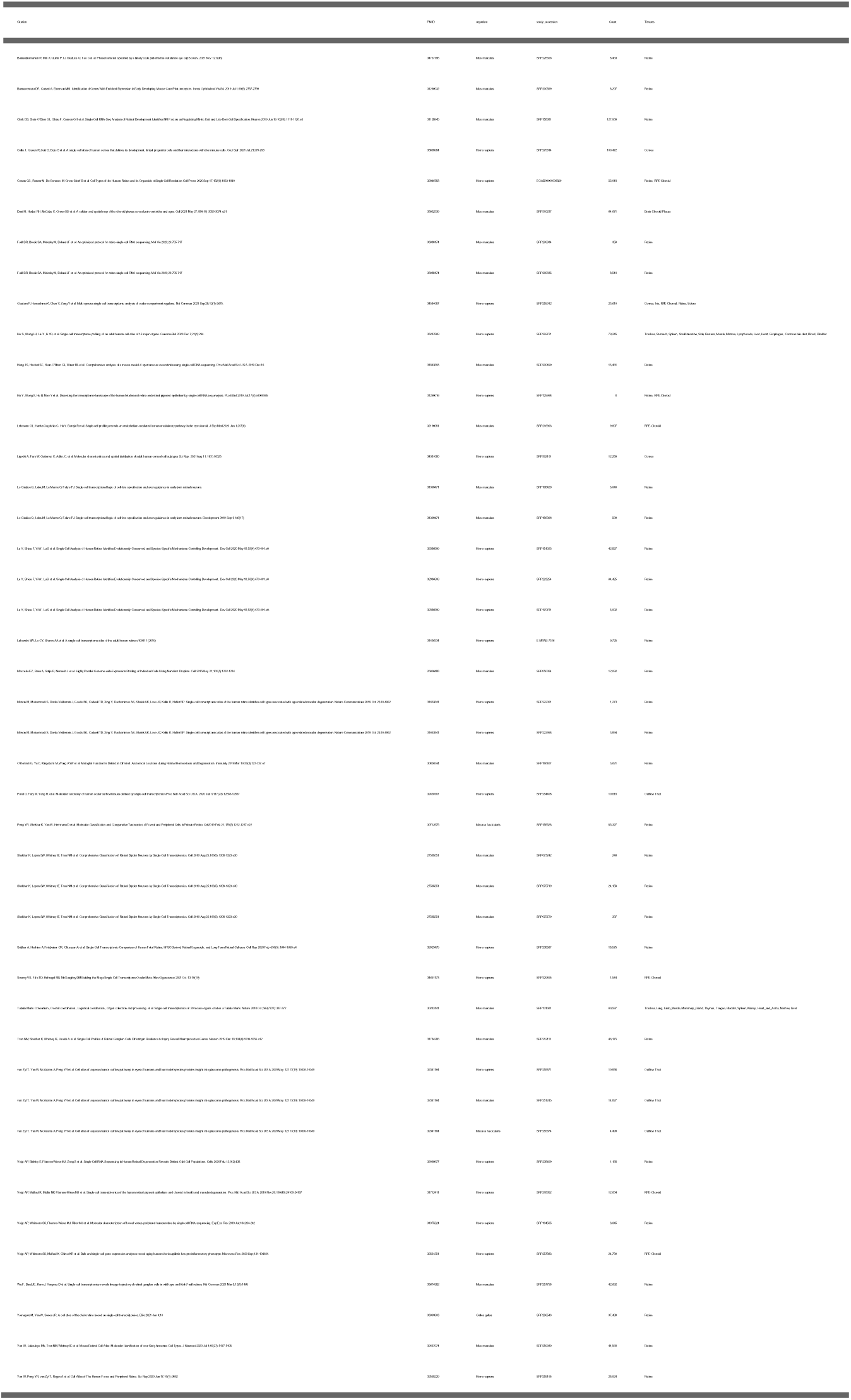
Studies, tissues, and number of cells across scEiaD

**Supplemental Table 2:**
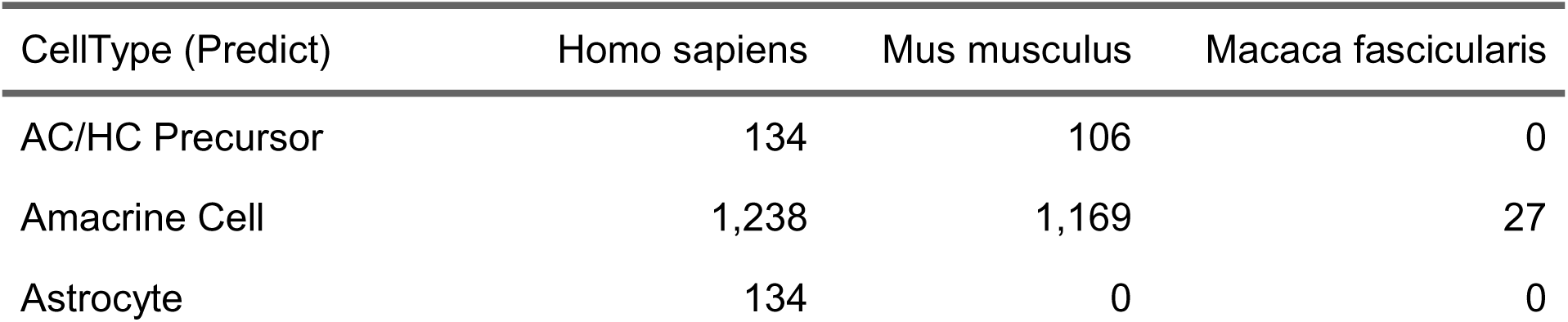
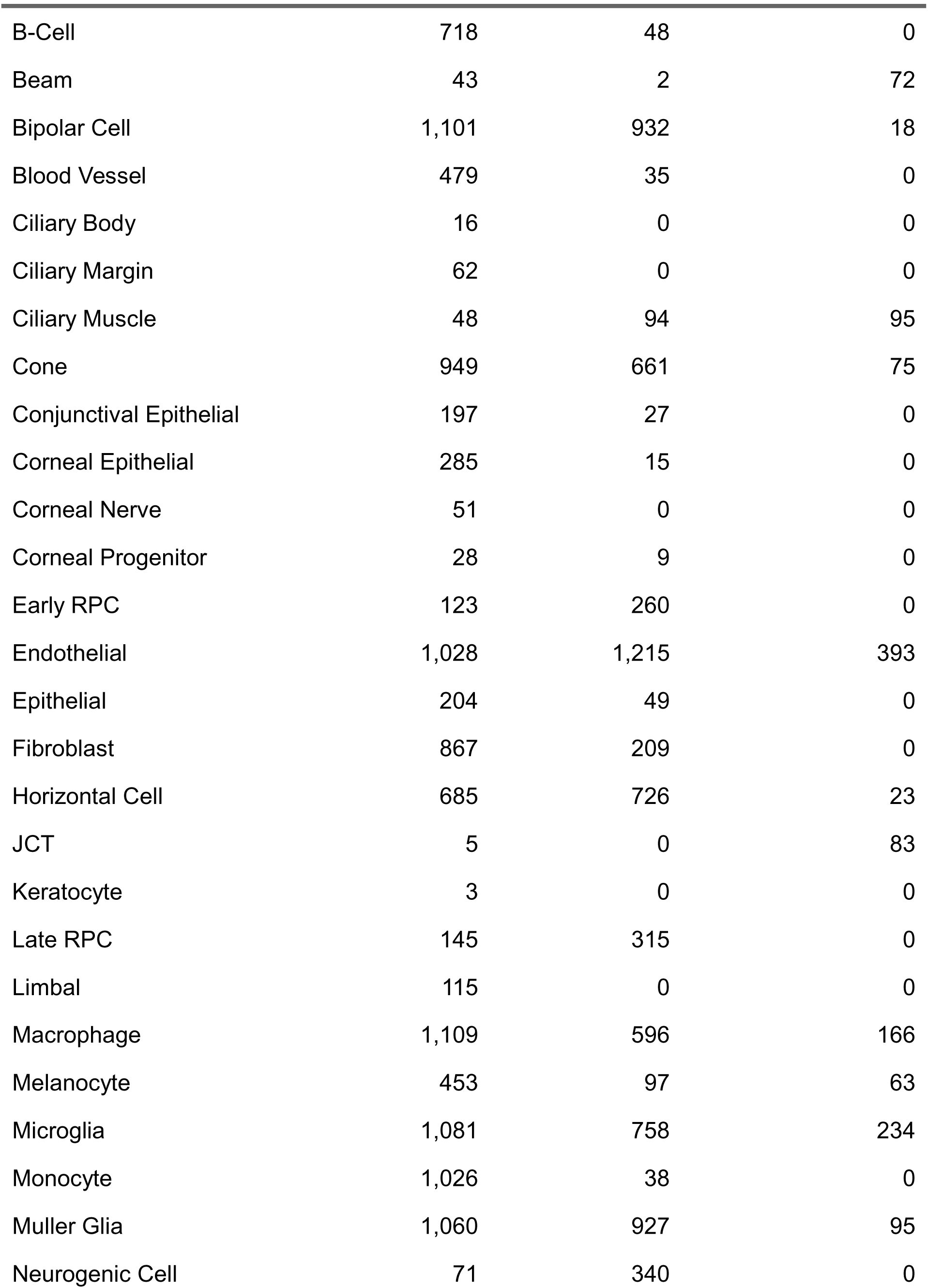
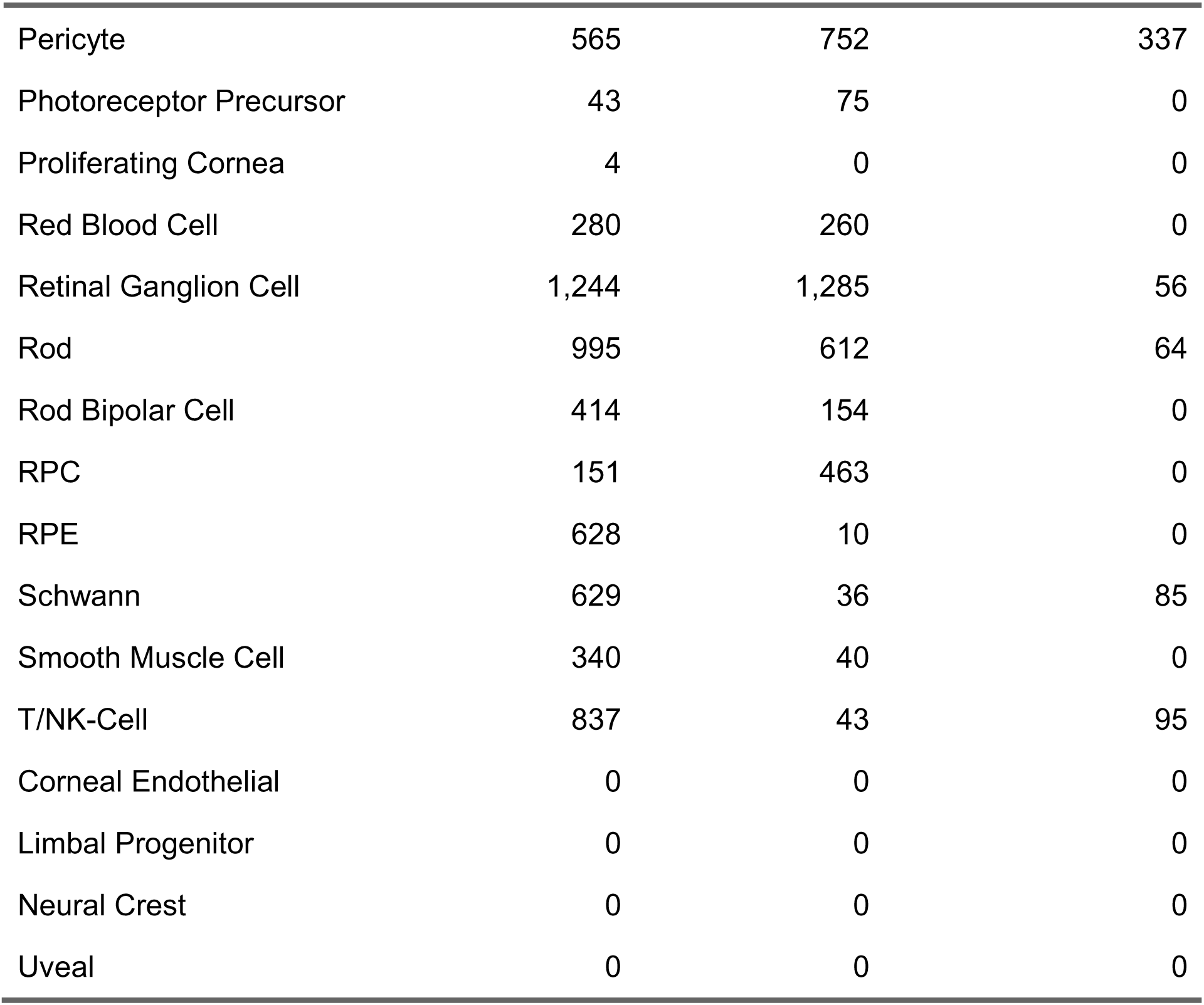
Counts of significantly differentially expressed genes for each cell type across the organisms

**Supplemental Table 3:** Counts of independent studies for each cell type across the organisms

| CellType (Predict) | Gallus gallus | Homo sapiens | Macaca fascicularis | Mus musculus | Well Supported |
| --- | --- | --- | --- | --- | --- |
| AC/HC Precursor | 0 | 3 | 0 | 3 | Yes |
| Amacrine Cell | 1 | 8 | 1 | 11 | Yes |
| Astrocyte | 0 | 2 | 0 | 0 | No |
| B-Cell | 0 | 5 | 0 | 0 | No |
| Beam | 0 | 3 | 1 | 1 | No |
| Bipolar Cell | 1 | 11 | 1 | 12 | Yes |
| Blood Vessel | 0 | 3 | 0 | 1 | No |
| Ciliary Body | 0 | 1 | 0 | 0 | No |
| Ciliary Margin | 0 | 1 | 0 | 1 | No |
| Ciliary Muscle | 0 | 2 | 1 | 1 | No |
| Cone | 1 | 8 | 1 | 9 | Yes |
| Conjunctival Epithelial | 0 | 3 | 0 | 1 | No |
| Corneal Endothelial | 0 | 2 | 0 | 0 | No |
| Corneal Epithelial | 0 | 4 | 0 | 1 | No |
| Corneal Nerve | 0 | 2 | 0 | 0 | No |
| Corneal Progenitor | 0 | 4 | 0 | 1 | No |
| Early RPC | 0 | 4 | 0 | 5 | Yes |
| Endothelial | 0 | 10 | 2 | 7 | Yes |
| Epithelial | 0 | 3 | 0 | 1 | No |
| Fibroblast | 0 | 10 | 0 | 3 | Yes |
| Horizontal Cell | 1 | 5 | 1 | 8 | Yes |
| JCT | 0 | 2 | 1 | 0 | No |
| Keratocyte | 0 | 2 | 0 | 0 | No |
| Late RPC | 0 | 3 | 0 | 4 | Yes |
| Limbal | 0 | 2 | 0 | 0 | No |
| Limbal Progenitor | 0 | 1 | 0 | 0 | No |
| Macrophage | 0 | 5 | 1 | 3 | Yes |
| Melanocyte | 0 | 8 | 1 | 2 | Yes |
| Microglia | 0 | 11 | 1 | 5 | Yes |
| Monocyte | 0 | 4 | 0 | 0 | No |
| Muller Glia | 1 | 12 | 1 | 6 | Yes |
| Neural Crest | 0 | 1 | 0 | 0 | No |
| Neurogenic Cell | 0 | 3 | 0 | 5 | Yes |
| Pericyte | 0 | 7 | 2 | 5 | Yes |
| Photoreceptor Precursor | 0 | 2 | 0 | 3 | Yes |
| Proliferating Cornea | 0 | 1 | 0 | 0 | No |
| Red Blood Cell | 0 | 2 | 0 | 1 | No |
| Retinal Ganglion Cell | 1 | 8 | 1 | 11 | Yes |
| Rod | 1 | 11 | 1 | 9 | Yes |
| Rod Bipolar Cell | 0 | 3 | 0 | 2 | Yes |
| RPC | 0 | 4 | 0 | 5 | Yes |
| RPE | 0 | 4 | 0 | 1 | No |
| Schwann | 0 | 5 | 1 | 1 | No |
| Smooth Muscle Cell | 0 | 2 | 0 | 1 | No |
| T/NK-Cell | 0 | 7 | 1 | 0 | No |
| Uveal | 0 | 1 | 0 | 1 | No |

**Supplemental Table 4:**
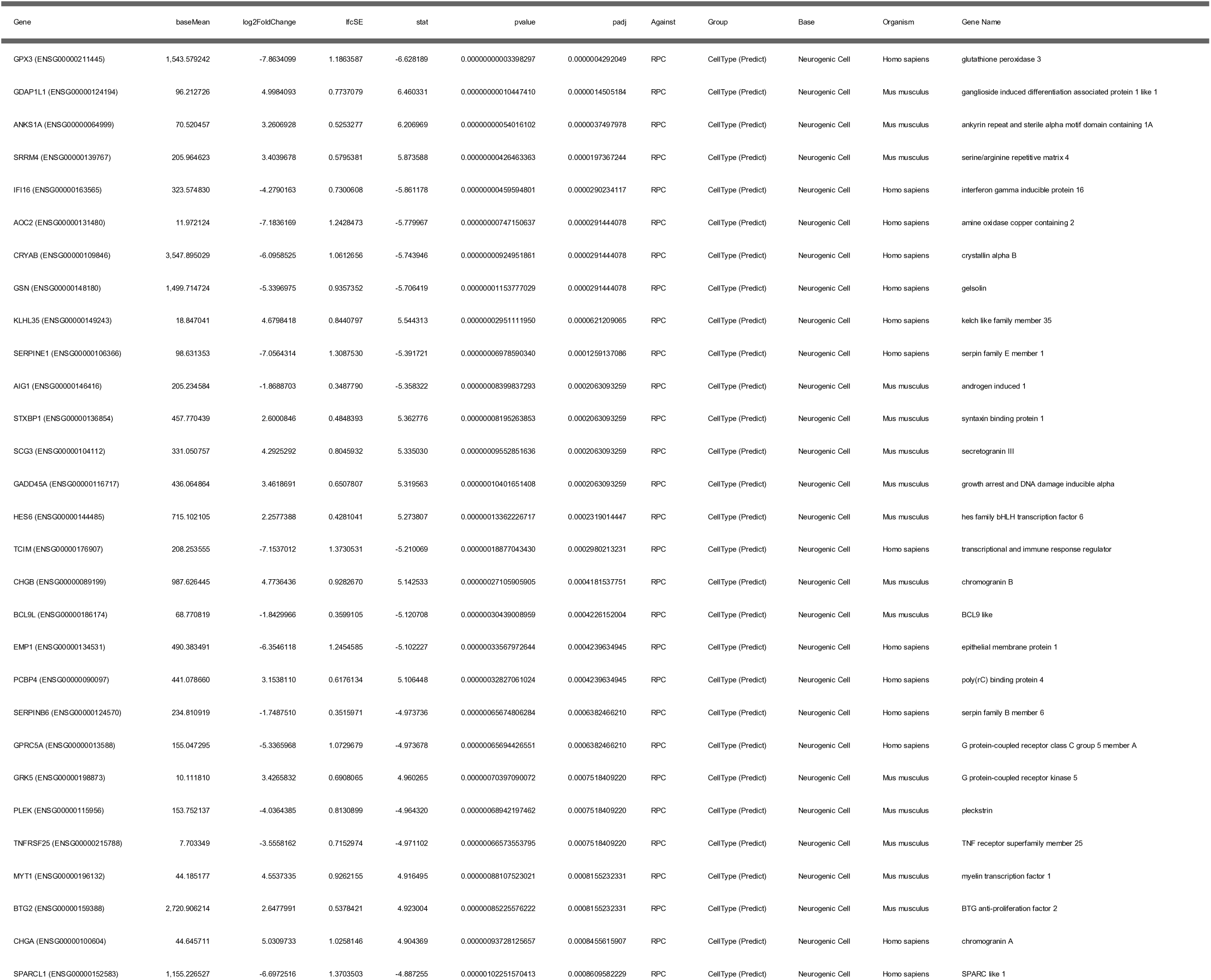
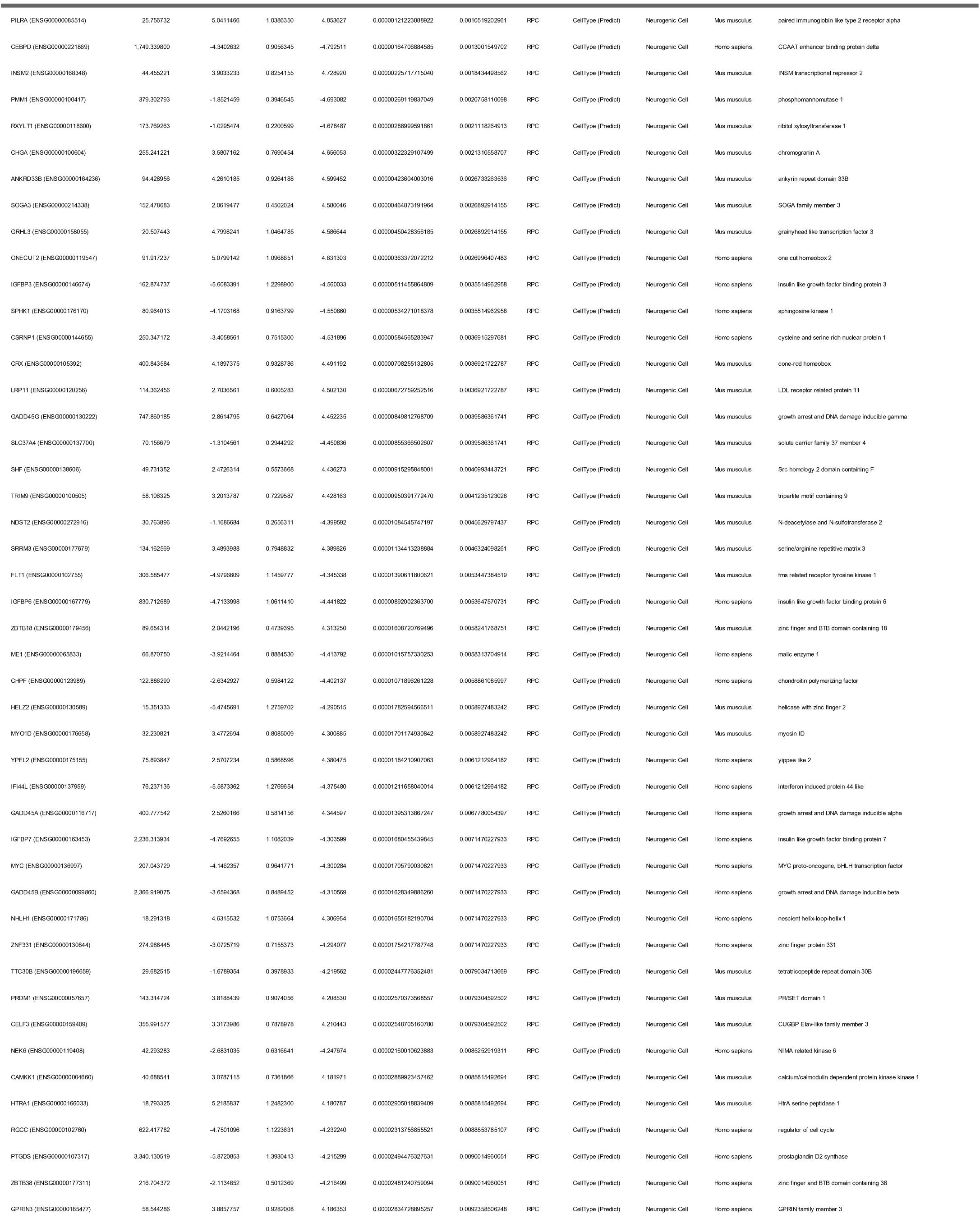
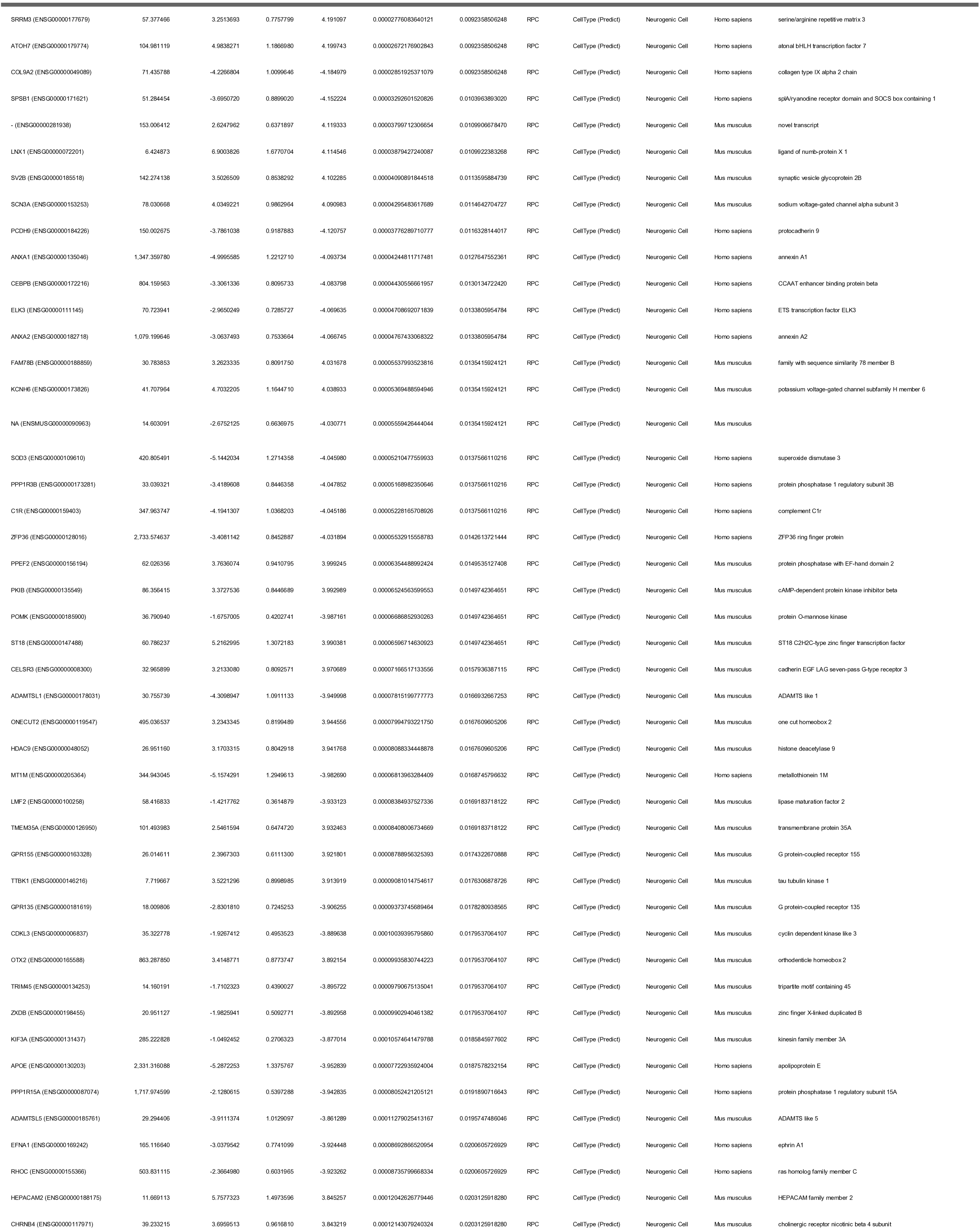
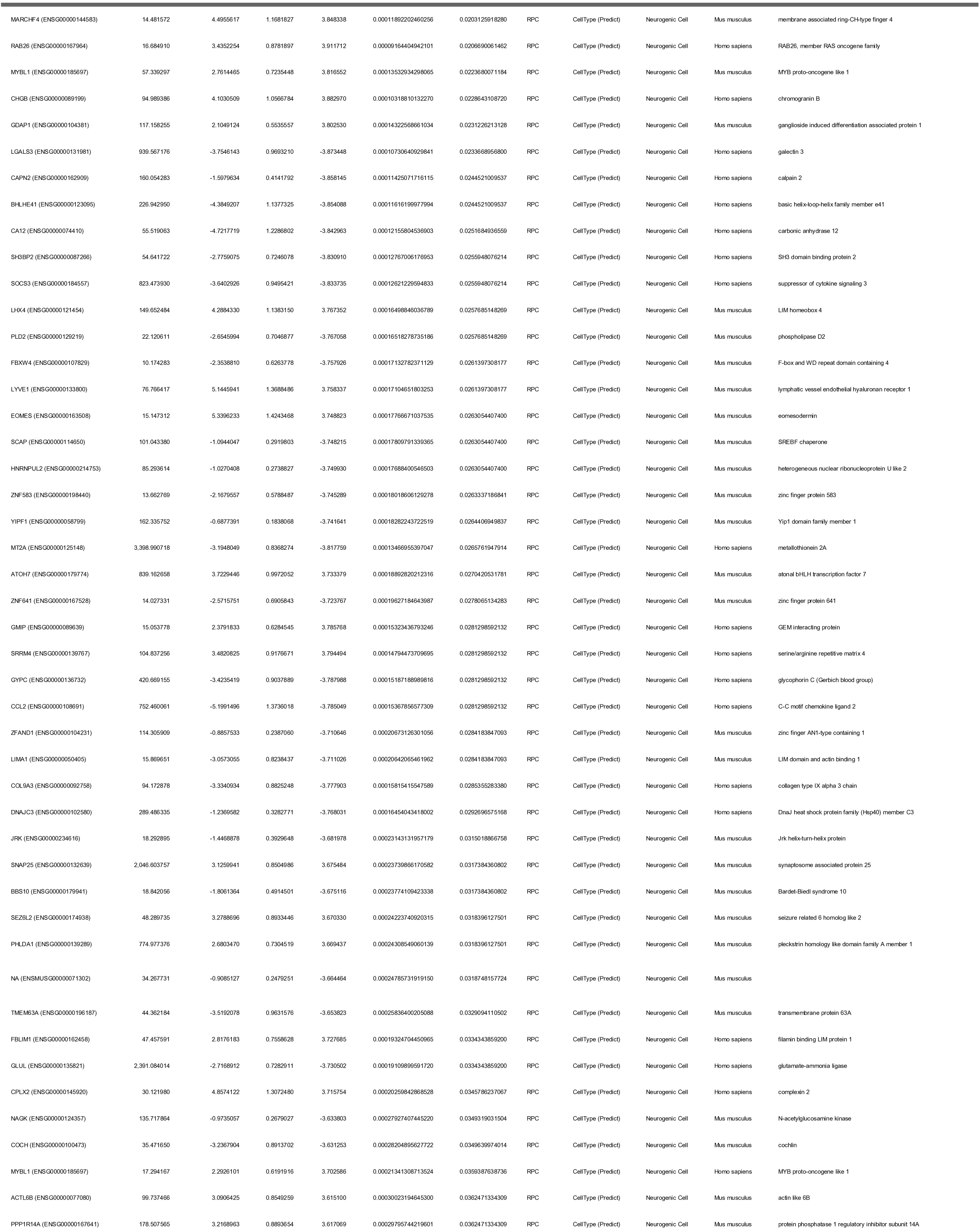
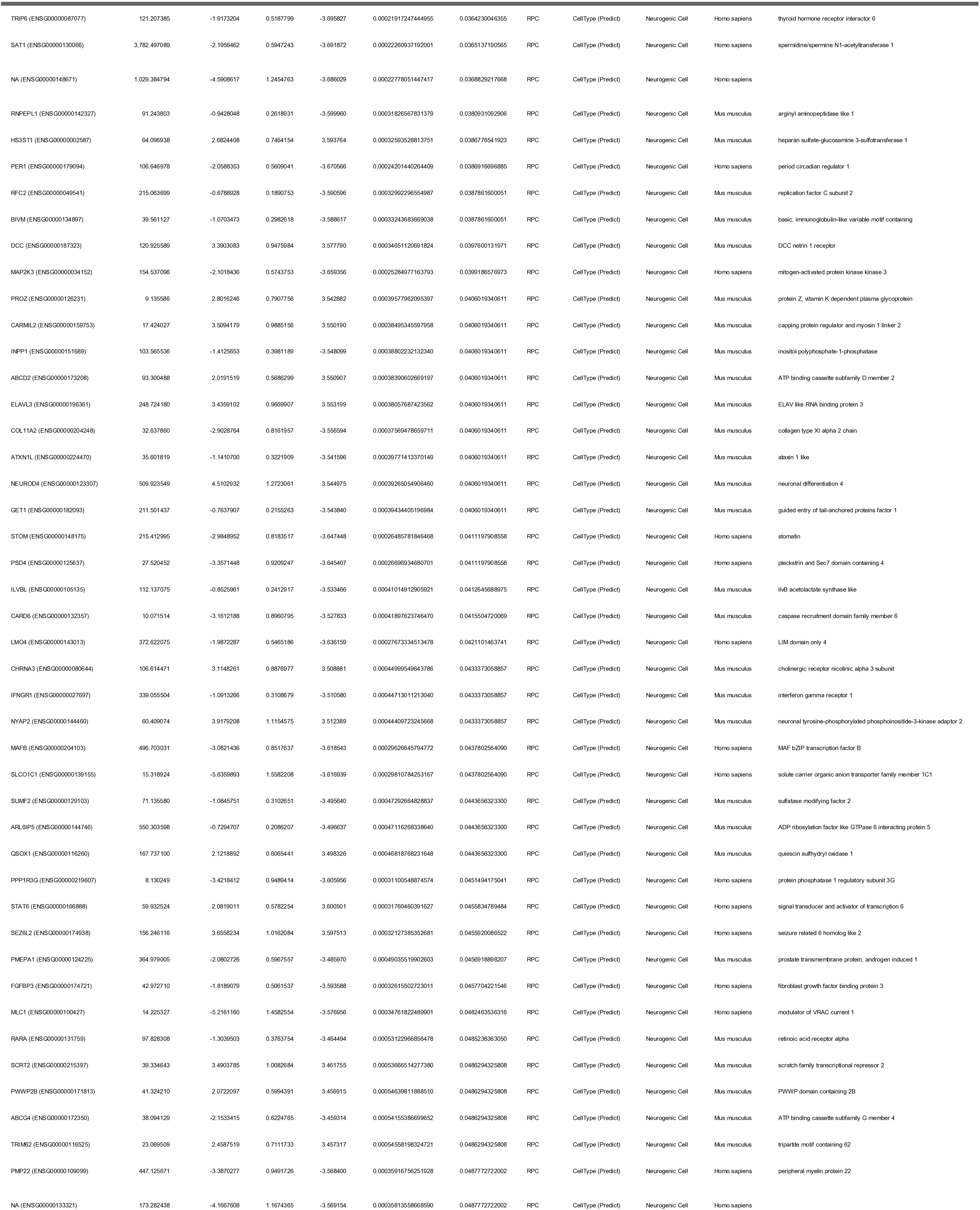
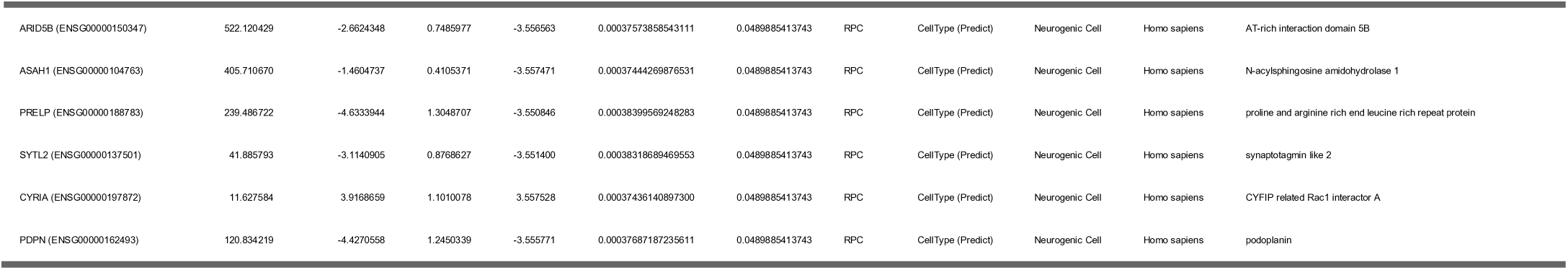
Differentially expressed genes between RPC and neurogenic cells in human and mouse. Genes filtered with a abs(log2FoldChange) > 2 and a padj < 0.01.

**Supplemental Table 5:** Macula versus peripheral cone differential gene testing table

| Gene | Ensembl | baseMean | log2FoldChange | lfcSE | stat | pvalue | padj |
| --- | --- | --- | --- | --- | --- | --- | --- |
| PDE6G | ENSG00000185527 | 66.333277 | -2.3441678 | 0.4168273 | -5.623835 | 0.00000001867637 | 0.0000490068 |
| AKAP12 | ENSG00000131016 | 16.211349 | 3.2204333 | 0.6509519 | 4.947268 | 0.00000075262380 | 0.0005739686 |
| HSPB1 | ENSG00000106211 | 33.905799 | 2.7358265 | 0.5443945 | 5.025449 | 0.00000050225706 | 0.0005739686 |
| GADD45B | ENSG00000099860 | 10.407572 | 2.7951199 | 0.5683610 | 4.917861 | 0.00000087495220 | 0.0005739686 |
| CC2D2A | ENSG00000048342 | 65.778406 | -1.8123582 | 0.4122057 | -4.396733 | 0.00001098925099 | 0.0057671589 |
| CALB1 | ENSG00000104327 | 18.454365 | 2.8703664 | 0.6662940 | 4.307958 | 0.00001647686745 | 0.0072058834 |
| RP1L1 | ENSG00000183638 | 27.760158 | -2.0706619 | 0.4881859 | -4.241544 | 0.00002219871789 | 0.0083213480 |
| RRAD | ENSG00000166592 | 10.848913 | 2.8671937 | 0.6933500 | 4.135276 | 0.00003545284876 | 0.0116285344 |
| H2AJ | ENSG00000246705 | 14.256520 | -1.7133388 | 0.4374781 | -3.916399 | 0.00008988147974 | 0.0262054448 |
| GNGT2 | ENSG00000167083 | 21.586045 | -1.8518370 | 0.4832323 | -3.832188 | 0.00012700851355 | 0.0333270340 |
| SLC24A4 | ENSG00000140090 | 12.091555 | -2.3479519 | 0.6275797 | -3.741281 | 0.00018308493710 | 0.0391603991 |
| EIF3E | ENSG00000104408 | 25.640591 | 1.7284801 | 0.4593680 | 3.762735 | 0.00016806515137 | 0.0391603991 |
| GPM6A | ENSG00000150625 | 14.342849 | 2.0169193 | 0.5412094 | 3.726690 | 0.00019401112370 | 0.0391603991 |
| SLC6A8 | ENSG00000130821 | 18.194596 | -1.5484910 | 0.4190325 | -3.695395 | 0.00021954477814 | 0.0411489641 |
| ATP8 | ENSG00000228253 | 24.808372 | -2.3805564 | 0.6505116 | -3.659514 | 0.00025269395502 | 0.0414418086 |
| JUN | ENSG00000177606 | 103.963097 | 2.0746674 | 0.5646982 | 3.673940 | 0.00023883868696 | 0.0414418086 |
| LBH | ENSG00000213626 | 22.178216 | -1.5820553 | 0.4438071 | -3.564736 | 0.00036422213745 | 0.0562187582 |
| MARCKS | ENSG00000277443 | 38.040849 | 1.4382002 | 0.4104345 | 3.504092 | 0.00045816776633 | 0.0632753799 |
| ATP8A2 | ENSG00000132932 | 15.822891 | -1.7063362 | 0.4867345 | -3.505681 | 0.00045543997512 | 0.0632753799 |
| SORL1 | ENSG00000137642 | 9.295426 | -2.0038012 | 0.5777338 | -3.468382 | 0.00052360288684 | 0.0654254274 |
| IGF1R | ENSG00000140443 | 10.280407 | 1.6259991 | 0.4675512 | 3.477692 | 0.00050575055876 | 0.0654254274 |
| PSIP1 | ENSG00000164985 | 10.390270 | 1.6926238 | 0.4991538 | 3.390987 | 0.00069641472693 | 0.0730080016 |
| AP1S2 | ENSG00000182287 | 15.468753 | 1.3309232 | 0.3949131 | 3.370167 | 0.00075122562584 | 0.0730080016 |
| PDLIM1 | ENSG00000107438 | 18.097895 | -1.6983405 | 0.5009208 | -3.390437 | 0.00069781251494 | 0.0730080016 |
| MCF2 | ENSG00000101977 | 7.817816 | -2.5509416 | 0.7532575 | -3.386546 | 0.00070778317976 | 0.0730080016 |
| ANKRD33 | ENSG00000167612 | 31.069622 | -1.6982079 | 0.4976874 | -3.412198 | 0.00064441375827 | 0.0730080016 |
| TRAC | ENSG00000277734 | 37.916861 | -1.4915835 | 0.4413536 | -3.379566 | 0.00072600465462 | 0.0730080016 |
| RIMS2 | ENSG00000176406 | 178.998521 | -0.9930176 | 0.2976702 | -3.335966 | 0.00085003661447 | 0.0796605742 |
| DNAJB1 | ENSG00000132002 | 44.874704 | 1.1598484 | 0.3504145 | 3.309933 | 0.00093318389388 | 0.0844370530 |
| FBL | ENSG00000105202 | 7.563700 | 1.9835304 | 0.6023149 | 3.293178 | 0.00099061590154 | 0.0866458709 |
| FOS | ENSG00000170345 | 65.413599 | 2.2598093 | 0.7217566 | 3.130985 | 0.00174220778280 | 0.0982445521 |
| UCKL1 | ENSG00000198276 | 21.260475 | -1.4652520 | 0.4673063 | -3.135528 | 0.00171544977959 | 0.0982445521 |
| DHRS11 | ENSG00000278535 | 27.726544 | -1.4295704 | 0.4514196 | -3.166832 | 0.00154109089340 | 0.0982445521 |
| PFKP | ENSG00000067057 | 9.676249 | -2.0719797 | 0.6594725 | -3.141874 | 0.00167870055349 | 0.0982445521 |
| NR1D2 | ENSG00000174738 | 15.146679 | 1.5612863 | 0.4908196 | 3.180978 | 0.00146778954795 | 0.0982445521 |
| ELOVL6 | ENSG00000170522 | 7.849669 | -1.8862844 | 0.5887784 | -3.203726 | 0.00135661747463 | 0.0982445521 |
| RAC1 | ENSG00000136238 | 17.670474 | 1.3239688 | 0.4090086 | 3.237019 | 0.00120785235883 | 0.0982445521 |
| BAMBI | ENSG00000095739 | 7.614820 | 2.6095010 | 0.8164728 | 3.196066 | 0.00139315244017 | 0.0982445521 |
| C12orf57 | ENSG00000111678 | 29.288427 | 1.0680875 | 0.3410052 | 3.132174 | 0.00173517129937 | 0.0982445521 |
| SGK1 | ENSG00000118515 | 15.018077 | -1.6407528 | 0.5122446 | -3.203065 | 0.00135973340152 | 0.0982445521 |
| KBTBD11 | ENSG00000176595 | 9.955960 | -1.8440108 | 0.5833912 | -3.160848 | 0.00157310538747 | 0.0982445521 |
| NSA2 | ENSG00000164346 | 13.607928 | 1.2706136 | 0.3961188 | 3.207658 | 0.00133820604225 | 0.0982445521 |
| RPL6 | ENSG00000089009 | 42.284446 | 1.1969586 | 0.3826535 | 3.128048 | 0.00175971568145 | 0.0982445521 |
| LIMA1 | ENSG00000050405 | 29.288181 | -1.4856845 | 0.4719500 | -3.147970 | 0.00164408359061 | 0.0982445521 |
| MCL1 | ENSG00000143384 | 11.135773 | 1.3862735 | 0.4282330 | 3.237195 | 0.00120711049749 | 0.0982445521 |
| TMEM14B | ENSG00000137210 | 20.558167 | -1.3397530 | 0.4279596 | -3.130559 | 0.00174473827329 | 0.0982445521 |
| USP32 | ENSG00000170832 | 17.570164 | -1.4198796 | 0.4481899 | -3.168031 | 0.00153474949030 | 0.0982445521 |

