## Supplementary material for "PLAE web app enables powerful searching and multiple visualizations across one million unified single-cell ocular transcriptomes": pseudobulk_cone_region_analysis

Pseudobulk Differential Testing of Fovea vs Peripheral Cones


Code 

- Show All Code
- Hide All Code
- Download Rmd

### Pseudobulk Differential Testing of Fovea vs Peripheral Cones


```
library(Seurat)
```


```
Registered S3 method overwritten by 'spatstat.geom':
  method     from
  print.boxx cli 
Attaching SeuratObject
```


```
library(tidyverse)
library(SingleCellExperiment)
```


```
Loading required package: SummarizedExperiment
Loading required package: MatrixGenerics
Loading required package: matrixStats

Attaching package: 'matrixStats'

The following object is masked from 'package:dplyr':

    count


Attaching package: 'MatrixGenerics'

The following objects are masked from 'package:matrixStats':

    colAlls, colAnyNAs, colAnys, colAvgsPerRowSet, colCollapse, colCounts, colCummaxs, colCummins, colCumprods, colCumsums, colDiffs, colIQRDiffs,
    colIQRs, colLogSumExps, colMadDiffs, colMads, colMaxs, colMeans2, colMedians, colMins, colOrderStats, colProds, colQuantiles, colRanges,
    colRanks, colSdDiffs, colSds, colSums2, colTabulates, colVarDiffs, colVars, colWeightedMads, colWeightedMeans, colWeightedMedians,
    colWeightedSds, colWeightedVars, rowAlls, rowAnyNAs, rowAnys, rowAvgsPerColSet, rowCollapse, rowCounts, rowCummaxs, rowCummins, rowCumprods,
    rowCumsums, rowDiffs, rowIQRDiffs, rowIQRs, rowLogSumExps, rowMadDiffs, rowMads, rowMaxs, rowMeans2, rowMedians, rowMins, rowOrderStats,
    rowProds, rowQuantiles, rowRanges, rowRanks, rowSdDiffs, rowSds, rowSums2, rowTabulates, rowVarDiffs, rowVars, rowWeightedMads,
    rowWeightedMeans, rowWeightedMedians, rowWeightedSds, rowWeightedVars

Loading required package: GenomicRanges
Loading required package: stats4
Loading required package: BiocGenerics

Attaching package: 'BiocGenerics'

The following object is masked from 'package:flextable':

    width

The following objects are masked from 'package:dplyr':

    combine, intersect, setdiff, union

The following objects are masked from 'package:stats':

    IQR, mad, sd, var, xtabs

The following objects are masked from 'package:base':

    anyDuplicated, append, as.data.frame, basename, cbind, colnames, dirname, do.call, duplicated, eval, evalq, Filter, Find, get, grep, grepl,
    intersect, is.unsorted, lapply, Map, mapply, match, mget, order, paste, pmax, pmax.int, pmin, pmin.int, Position, rank, rbind, Reduce,
    rownames, sapply, setdiff, sort, table, tapply, union, unique, unsplit, which.max, which.min

Loading required package: S4Vectors

Attaching package: 'S4Vectors'

The following objects are masked from 'package:dplyr':

    first, rename

The following object is masked from 'package:tidyr':

    expand

The following objects are masked from 'package:base':

    expand.grid, I, unname

Loading required package: IRanges

Attaching package: 'IRanges'

The following object is masked from 'package:glue':

    trim

The following objects are masked from 'package:dplyr':

    collapse, desc, slice

The following object is masked from 'package:purrr':

    reduce

Loading required package: GenomeInfoDb
Loading required package: Biobase
Welcome to Bioconductor

    Vignettes contain introductory material; view with 'browseVignettes()'. To cite Bioconductor, see 'citation("Biobase")', and for packages
    'citation("pkgname")'.


Attaching package: 'Biobase'

The following object is masked from 'package:MatrixGenerics':

    rowMedians

The following objects are masked from 'package:matrixStats':

    anyMissing, rowMedians


Attaching package: 'SummarizedExperiment'

The following object is masked from 'package:SeuratObject':

    Assays

The following object is masked from 'package:Seurat':

    Assays
```


```
library(scran)
```


```
Loading required package: scuttle
```


```
library(org.Hs.eg.db)
```


```
Loading required package: AnnotationDbi

Attaching package: 'AnnotationDbi'

The following object is masked from 'package:dplyr':

    select
```


```
#system('wget -O ~/data/scEiaD_2022_02/scEiaD_all_seurat_v3.Rdata https://hpc.nih.gov/~mcgaugheyd/scEiaD/2022_03_22/scEiaD_all_seurat_v3.Rdata)
# the -O renames the file and puts it in the ~/data/scEiaD directory
load('~/data/scEiaD_2022_02/scEiaD_all_seurat_v3.Rdata')
```


#### Compare macula cones vs peripheral cones

Let us now demonstrate how having a large annotated set of data can be of value in asking more specific questions, like “how do macula/fovea cones differ relative to peripheral cones.”

First, we need to see whether this is even a tractable question.


```
scEiaD@meta.data %>% 
  as_tibble() %>% 
  group_by(retina_region, organism, study_accession, CellType_predict) %>% 
  summarise(Count = n()) %>% 
  filter(CellType_predict %in% c("Cone")) %>% 
  filter(Count > 10, !is.na(retina_region))
```


```
`summarise()` has grouped output by 'retina_region', 'organism', 'study_accession'. You can override using the `.groups` argument.
```


Yes, we have several human studies with macula / peripheral cone cells.

Let’s demonstrate how we can quickly run a conservative differential expression test the leverages the many studies we have. The “traditional” scRNA diff tests often return insane numbers of highly differentially expressed genes (the test we ran aboves returns many many genes with a padj < 0.05), which is highly annoying as then you really have to plot each result individually to confirm that it does not look wacky. We are going to build a “pseudo bulk” experiment here where we sum all the counts by gene, cell type (in this case we are doing macula cones *versus* peripheral cones), and study. This “pseudo” count table can then be used in a classic differential testing system like edgeR, limma, or DESeq2 (we will use the later).


```
# Create new column with a pasted together retina region and a CellType
scEiaD <- AddMetaData(scEiaD, 
                      metadata = 
                        paste0(
                          scEiaD@meta.data$retina_region, 
                          '_', 
                          scEiaD@meta.data$CellType_predict,
                          '_',
                          scEiaD@meta.data$organism
                        ), 
                      col.name = 'regionCT')


# filter down scEiaD to only human and in semi-supported (more than 10 cones cells) studies
Idents(scEiaD) <- scEiaD@meta.data$regionCT
scEiaD__subset <- subset(scEiaD, idents = c('Macula_Cone_Homo sapiens','Peripheral_Cone_Homo sapiens'))
```


##### Wrinkle

scEiaD is built on the Ensembl gene id (for examples ENSG00000185527). That’s not very friendly to humans so I added a bit of code to link the Ensembl ID to the gene name and show the top 16 diff genes (padj < 0.05) as a table.


```
# get human gene names 
library(org.Hs.eg.db)
symbols <- mapIds(org.Hs.eg.db, keys = row.names(deseq_res), keytype = "ENSEMBL", column="SYMBOL")
```


```
'select()' returned 1:many mapping between keys and columns
```


```
region_table <- deseq_res %>% as_tibble(rownames = 'name') %>%  
  left_join(symbols %>% enframe()) %>% 
  dplyr::rename(Ensembl = name, Gene = value) %>% 
  relocate(Gene) %>% 
  filter(padj < 0.05)
```


```
Joining, by = "name"
```


```
region_table %>% 
  arrange(pvalue) %>% DT::datatable()
```

##### Always look at the plots

We use the “ScaleData” function from Seurat to zero center and scale the counts for each dataset #### PDE6G Top gene more highly expressed in the macula compared to the periphery. Seems pretty consistently higher across all the studies.


```
scEiaD__subset <- ScaleData(scEiaD__subset)
```


```
Centering and scaling data matrix

  |                                                                                                                                                                
  |                                                                                                                                                          |   0%
  |                                                                                                                                                                
  |======================================                                                                                                                    |  25%
  |                                                                                                                                                                
  |=============================================================================                                                                             |  50%
  |                                                                                                                                                                
  |====================================================================================================================                                      |  75%
  |                                                                                                                                                                
  |==========================================================================================================================================================| 100%
```


```
VlnPlot(scEiaD__subset, features = c('ENSG00000185527'), log = TRUE)
```


```
VlnPlot(scEiaD__subset, c('ENSG00000185527'), split.by = 'retina_region', group.by='study_accession', log = TRUE)
```


```
The default behaviour of split.by has changed.
Separate violin plots are now plotted side-by-side.
To restore the old behaviour of a single split violin,
set split.plot = TRUE.
      
This message will be shown once per session.
```


###### All genes


```
for (i in region_table$Ensembl){
  #print(i)
  print(VlnPlot(scEiaD__subset, i, split.by = 'retina_region', group.by='study_accession', log = TRUE) )
}
```

### Output and Session Info


```
sessionInfo()
```


```
R version 4.1.2 (2021-11-01)
Platform: x86_64-apple-darwin17.0 (64-bit)
Running under: macOS Catalina 10.15.7

Matrix products: default
BLAS:   /System/Library/Frameworks/Accelerate.framework/Versions/A/Frameworks/vecLib.framework/Versions/A/libBLAS.dylib
LAPACK: /Library/Frameworks/R.framework/Versions/4.1/Resources/lib/libRlapack.dylib

locale:
[1] en_US.UTF-8/en_US.UTF-8/en_US.UTF-8/C/en_US.UTF-8/en_US.UTF-8

attached base packages:
[1] stats4    stats     graphics  grDevices utils     datasets  methods   base     

other attached packages:
 [1] DESeq2_1.34.0               org.Hs.eg.db_3.14.0         AnnotationDbi_1.56.1        scran_1.22.1                scuttle_1.4.0              
 [6] SingleCellExperiment_1.16.0 SummarizedExperiment_1.24.0 Biobase_2.54.0              GenomicRanges_1.46.0        GenomeInfoDb_1.30.0        
[11] IRanges_2.28.0              S4Vectors_0.32.2            BiocGenerics_0.40.0         MatrixGenerics_1.6.0        matrixStats_0.61.0         
[16] SeuratObject_4.0.4          Seurat_4.0.6                shiny_1.7.1                 RSQLite_2.2.8               pool_0.1.6                 
[21] tictoc_1.0.1                glue_1.5.0                  captioner_2.2.3             flextable_0.6.10            colorspace_2.0-2           
[26] ggrepel_0.9.1               cowplot_1.1.1               citr_0.3.2                  forcats_0.5.1               stringr_1.4.0              
[31] dplyr_1.0.7                 purrr_0.3.4                 readr_2.0.2                 tidyr_1.1.4                 tibble_3.1.6               
[36] ggplot2_3.3.5               tidyverse_1.3.1            

loaded via a namespace (and not attached):
  [1] rappdirs_0.3.3            rtracklayer_1.54.0        scattermore_0.7           bit64_4.0.5               knitr_1.36                fst_0.9.4                
  [7] irlba_2.3.5               DelayedArray_0.20.0       rpart_4.1-15              data.table_1.14.2         KEGGREST_1.34.0           RCurl_1.98-1.5           
 [13] AnnotationFilter_1.18.0   doParallel_1.0.16         generics_0.1.1            GenomicFeatures_1.46.5    ScaledMatrix_1.2.0        RANN_2.6.1               
 [19] future_1.23.0             bit_4.0.4                 tzdb_0.2.0                spatstat.data_2.1-2       xml2_1.3.2                lubridate_1.8.0          
 [25] httpuv_1.6.3              assertthat_0.2.1          fontawesome_0.2.2         viridis_0.6.2             xfun_0.28                 hms_1.1.1                
 [31] jquerylib_0.1.4           evaluate_0.14             promises_1.2.0.1          fansi_0.5.0               restfulr_0.0.13           progress_1.2.2           
 [37] dbplyr_2.1.1              readxl_1.3.1              geneplotter_1.72.0        igraph_1.2.8              DBI_1.1.1                 htmlwidgets_1.5.4        
 [43] spatstat.geom_2.3-1       ellipsis_0.3.2            crosstalk_1.2.0           backports_1.3.0           annotate_1.72.0           deldir_1.0-6             
 [49] biomaRt_2.50.0            sparseMatrixStats_1.6.0   vctrs_0.3.8               ensembldb_2.18.4          ROCR_1.0-11               abind_1.4-5              
 [55] cachem_1.0.6              withr_2.4.2               ggforce_0.3.3             sctransform_0.3.3         GenomicAlignments_1.30.0  prettyunits_1.1.1        
 [61] goftest_1.2-3             cluster_2.1.2             lazyeval_0.2.2            crayon_1.4.2              genefilter_1.76.0         edgeR_3.36.0             
 [67] pkgconfig_2.0.3           labeling_0.4.2            tweenr_1.0.2              nlme_3.1-153              vipor_0.4.5               ProtGenerics_1.26.0      
 [73] pals_1.7                  rlang_0.4.12              globals_0.14.0            lifecycle_1.0.1           miniUI_0.1.1.1            filelock_1.0.2           
 [79] BiocFileCache_2.2.0       modelr_0.1.8              rsvd_1.0.5                dichromat_2.0-0           cellranger_1.1.0          polyclip_1.10-0          
 [85] lmtest_0.9-39             Matrix_1.3-4              zoo_1.8-9                 reprex_2.0.1              base64enc_0.1-3           beeswarm_0.4.0           
 [91] ggridges_0.5.3            GlobalOptions_0.1.2       png_0.1-7                 viridisLite_0.4.0         rjson_0.2.20              bitops_1.0-7             
 [97] KernSmooth_2.23-20        Biostrings_2.62.0         blob_1.2.2                DelayedMatrixStats_1.16.0 shape_1.4.6               parallelly_1.30.0        
[103] beachmat_2.10.0           scales_1.1.1              memoise_2.0.0             magrittr_2.0.1            plyr_1.8.6                ica_1.0-2                
[109] bibtex_0.4.2.3            zlibbioc_1.40.0           compiler_4.1.2            RefManageR_1.3.0          dqrng_0.3.0               BiocIO_1.4.0             
[115] RColorBrewer_1.1-2        clue_0.3-60               fitdistrplus_1.1-6        Rsamtools_2.10.0          cli_3.1.0                 XVector_0.34.0           
[121] listenv_0.8.0             pbapply_1.5-0             patchwork_1.1.1           mgcv_1.8-38               MASS_7.3-54               tidyselect_1.1.1         
[127] stringi_1.7.5             yaml_2.2.1                locfit_1.5-9.4            BiocSingular_1.10.0       grid_4.1.2                sass_0.4.0               
[133] tools_4.1.2               future.apply_1.8.1        parallel_4.1.2            circlize_0.4.13           rstudioapi_0.13           uuid_1.0-3               
[139] bluster_1.4.0             foreach_1.5.1             metapod_1.2.0             gridExtra_2.3             farver_2.1.0              Rtsne_0.15               
[145] digest_0.6.28             Rcpp_1.0.7                broom_0.7.10              later_1.3.0               RcppAnnoy_0.0.19          httr_1.4.3               
[151] gdtools_0.2.3             ComplexHeatmap_2.10.0     tensor_1.5                rvest_1.0.2               XML_3.99-0.8              fs_1.5.0                 
[157] reticulate_1.22           splines_4.1.2             statmod_1.4.36            uwot_0.1.11               spatstat.utils_2.3-0      scater_1.22.0            
[163] mapproj_1.2.7             plotly_4.10.0             systemfonts_1.0.2         xtable_1.8-4              jsonlite_1.7.2            R6_2.5.1                 
[169] pillar_1.6.4              htmltools_0.5.2           mime_0.12                 fastmap_1.1.0             DT_0.19                   BiocParallel_1.28.0      
[175] BiocNeighbors_1.12.0      codetools_0.2-18          maps_3.4.0                utf8_1.2.2                spatstat.sparse_2.1-0     lattice_0.20-45          
[181] bslib_0.3.1               curl_4.3.2                ggbeeswarm_0.6.0          leiden_0.3.9              officer_0.4.1             zip_2.2.0                
[187] shinyjs_2.0.0             limma_3.50.0              survival_3.2-13           rmarkdown_2.11            munsell_0.5.0             GetoptLong_1.0.5         
[193] GenomeInfoDbData_1.2.7    iterators_1.0.13          reshape2_1.4.4            haven_2.4.3               gtable_0.3.0              spatstat.core_2.3-2
```


```
save(scEiaD__subset,region_table, deseq_res, file = 'pseudoBulk_cone_region_files.Rdata' )
write_csv(region_table, file = 'pseudoBulk_cone_region_table.csv.gz' )
```
