## Supplementary material for "PLAE web app enables powerful searching and multiple visualizations across one million unified single-cell ocular transcriptomes": R scripts: diff_data_pull.R

```
library(tidyverse)

library(cowplot)
library(ggrepel)


library(glue)
library(tictoc)

tic()
#data_dir_vGiga <- '/Volumes/McGaughey_S/data/scEiaD//'
#data_dir_vPLAE <- '/Volumes/McGaughey_S/data/scEiaD_2022_02/'
data_dir_vGiga <- '~/data/scEiaD_gigascience/'
data_dir_vPLAE <- '~/data/scEiaD_2022_02/'


library(pool)
library(RSQLite)
meta_filter_v0 <- fst::read_fst(glue("{data_dir_vGiga}/paper_data/2021_03_17_meta_filter.fst")) %>% 
  
  mutate(CellType_predict = case_when(CellType_predict == 'AC/HC_Precurs' ~ 'AC/HC Precursor',
                                      TRUE ~ CellType_predict),
         CellType= case_when(CellType == 'AC/HC_Precurs' ~ 'AC/HC Precursor',
                             TRUE ~ CellType))

meta_filter <- fst::read_fst(glue("{data_dir_vPLAE}/meta_filter.fst")) %>% filter(study_accession != 'Bharti_Nguyen_iRPE_2D_3D') %>% 
  
  mutate(CellType_predict = case_when(CellType_predict == 'AC/HC_Precursor' ~ 'AC/HC Precursor',
                                      TRUE ~ CellType_predict),
         CellType= case_when(CellType == 'AC/HC_Precursor' ~ 'AC/HC Precursor',
                             TRUE ~ CellType),  
         Tissue = case_when(grepl('Iris', Tissue) ~ 'Iris',
                            TRUE ~ Tissue),
         Tissue = case_when(grepl('^RPE|^Choroid', Tissue) ~ 'RPE-Choroid',
                            TRUE ~ Tissue),
         Citation = case_when(study_accession == 'OGVFB_Hufnagel_iPSC_RPE' ~ 'Swamy VS, Fufa TD, Hufnagel RB, McGaughey DM Building the Mega Single Cell Transcriptome Ocular Meta-Atlas Gigascience 2021 Oct 13;10(10)',
                              TRUE ~ Citation),
         PMID = case_when(study_accession == 'OGVFB_Hufnagel_iPSC_RPE' ~ '34651173',
                          TRUE ~ PMID),
         study_accession = case_when(study_accession == 'OGVFB_Hufnagel_iPSC_RPE' ~ 'SRP329495',
                                     TRUE ~ study_accession))

scEiaD_2020_v00 <- pool::dbPool(RSQLite::SQLite(), dbname =  glue('{data_dir_vGiga}/MOARTABLES__anthology_limmaFALSE___5000-transform-counts-universe-batch-scVIprojectionSO-8-0.1-50-5.sqlite'))
scEiaD_2020_v01 <- pool::dbPool(RSQLite::SQLite(), dbname =  glue('{data_dir_vPLAE}/MOARTABLES__anthology_limmaFALSE___4000-counts-universe-study_accession-scANVIprojection-15-5-0.1-50-20__pointRelease01.sqlite'))

toc()


######################################


tic()
# number of celltype (predict) - organism - study counts (with >=50 cells)
# ws == Well Supported
ws_table <- meta_filter %>%  
  filter(Organ == 'Eye') %>% 
  group_by(organism, CellType_predict, study_accession) %>% 
  count() %>% 
  filter(n >= 50) %>% 
  ungroup() %>% 
  group_by(organism, CellType_predict) %>% 
  count() %>% 
  pivot_wider(values_from = n, names_from = organism) %>% 
  filter(!is.na(CellType_predict)) %>% 
  replace(is.na(.), values = 0) %>% 
  arrange(CellType_predict) %>% 
  rename(`CellType (Predict)` = CellType_predict) %>% 
  mutate(`Well Supported` = case_when(`Homo sapiens` >= 2 & `Mus musculus` >= 2 ~ 'Yes',
                                      TRUE ~ 'No'))

# fix  occasional NA
# data made from src/ensdb.R
diff_tab_big <- read_tsv('data/diff_tab_big.tsv.gz')
down_tab_big <- read_tsv('data/down_tab_big.tsv.gz')
gene_tab <- bind_rows(read_tsv('data/hs.ensdb.v86.symbol.geneid.tsv.gz'), read_tsv('data/mm.ensdb.v79.symbol.geneid.tsv.gz'))
diff_tab <- diff_tab_big %>% 
  filter(Against == 'All', 
         Base %in% (meta_filter %>% filter(Organ == 'Eye') %>% 
                      pull(CellType) %>% unique()))


diff_tab_sup <- diff_tab %>% 
  group_by(Base, Organism) %>% 
  summarise(Count = n()) %>% 
  rename(`CellType (Predict)` = Base, `Differentially Expressed Gene Count` = Count) %>% 
  pivot_wider(values_from = `Differentially Expressed Gene Count`, names_from = Organism) %>% replace(is.na(.),0) %>% 
  as_tibble()

diff_tab_sup <- bind_rows(
  diff_tab_sup,
  ws_table$`CellType (Predict)`[!ws_table$`CellType (Predict)` %in% 
                                  diff_tab_sup$`CellType (Predict)`] %>% 
    enframe() %>% 
    select(`CellType (Predict)` = value) %>%
    mutate(`Homo sapiens` = 0, `Mus musculus` = 0, `Macaca fascicularis` = 0) %>% 
    arrange(`CellType (Predict)`)
)

write_csv(diff_tab, file = 'supplemental_files/differential_expression_table.csv.gz')

# diff_tab_sup %>% 
#   flextable()
# 
# ws_table %>% 
#   flextable()

# diff_tab_barplot <- diff_tab_sup %>% 
#   pivot_longer(-`CellType (Predict)`, names_to = 'Organism', values_to = 'Diff Gene Count') %>% 
#   ggplot(aes(x=`CellType (Predict)`, y = `Diff Gene Count`, fill = Organism )) + 
#   geom_bar(stat='identity', position = 'dodge') + 
#   cowplot::theme_cowplot() + coord_flip()

toc()


#####################################
tic()
ws_oct <- ws_table %>% filter(`Well Supported` == 'Yes') %>% pull(`CellType (Predict)`)

consist_diff <- scEiaD_2020_v01 %>%
  tbl("diff_testing") %>%
  filter(Base %in% ws_oct) %>%
  filter(Group == 'CellType (Predict)',
         Against == 'All',
         Organism %in% c('Homo sapiens','Mus musculus')) %>%
  group_by(Base, Gene) %>%
  summarise(`mean log2FC` = mean(log2FoldChange), `sig count` = sum(padj < 1e-5), `mean padj` = mean(padj)) %>%
  filter(`mean log2FC` > 2, `sig count` > 1, `mean padj` < 1e-5) %>% 
  collect() %>%
  mutate(GENEID = str_extract(Gene, 'ENS\\w+')) %>%
  left_join(gene_tab, by = 'GENEID') %>%
  relocate(SYMBOL, GENEID) %>%
  dplyr::rename(Symbol = SYMBOL, ID = GENEID) %>%
  arrange(-`mean log2FC`)

# set genes to one cell types (if there are multiple, then pick the one with the highest mean log2FC)
# remove genes that are diff expressed in more than 3 diff cell types
top_genes <- consist_diff %>% group_by(Symbol) %>% summarise(Count = n()) %>% filter(Count <= 3) %>% left_join(consist_diff %>%  group_by(Symbol) %>% slice_max(., n=1, order_by = `mean log2FC`), by = 'Symbol')
# consist_diff %>% group_by(Base) %>% summarise(Count = n())
# consist_diff %>% filter(Symbol %in% uniq_g$Symbol) %>% group_by(Base) %>% summarise(Count = n())

# consist_diff %>% filter(Symbol %in% uniq_g$Symbol) %>% group_by(Base) %>% summarise(Count = n()) %>% ggplot(aes(x=`Base`, y = `Count` )) + geom_bar(stat='identity', position = 'dodge') + cowplot::theme_cowplot() + coord_flip() + xlab("CellTye (Predict)")
toc()
#######################################

tic()
g <- top_genes %>% select(Symbol, Base) %>% filter(!Base %in% c('Early RPC','Late RPC')) %>% left_join(consist_diff, by = c("Symbol", "Base"))  %>% group_by(Base) %>% slice_max(n=8, order_by = `mean log2FC`) %>% mutate(g = paste0(Symbol, ' (', ID, ')')) %>% pull(g)

consist_diff_long <- scEiaD_2020_v01 %>%
  tbl("diff_testing") %>%
  filter(Gene %in% g) %>%
  collect() %>%
  filter(Base %in% top_genes$Base,
         !Base %in% c('Early RPC','Late RPC'), 
         Group == 'CellType (Predict)',
         Against == 'All') %>%
  select(Gene, Organism, log2FoldChange, Base)

consist_diff_wide <- consist_diff_long %>% 
  mutate(GO = paste0(Base, '_', Organism)) %>% 
  select(-Organism, -Base) %>% 
  pivot_wider(values_from=log2FoldChange, names_from = GO)
consist_diff_celltype <- colnames(consist_diff_wide[,-1])
consist_diff_wide <- consist_diff_wide %>% data.frame()
row.names(consist_diff_wide) <- consist_diff_wide$Gene

consist_diff_wide <- consist_diff_wide[,-1]


consist_diff_organism <- str_extract(consist_diff_celltype, '_\\w+ \\w+') %>% gsub('_','',.)
consist_diff_cts  <- str_extract(consist_diff_celltype, '.*_') %>% gsub('_','',.)
consist_diff_genes <- row.names(consist_diff_wide) %>% gsub(' \\(.*','',.)

consist_diff_base <- consist_diff_genes %>% as_tibble() %>% rename(Symbol = value) %>% left_join(top_genes %>% select(Symbol, Base) %>% unique()) %>% pull(Base)

# 
# consist_diff_hm <- ComplexHeatmap::Heatmap(consist_diff_wide, column_labels = consist_diff_organism, 
#                                            column_split = consist_diff_cts, 
#                                            row_split = consist_diff_base, show_row_dend = FALSE, 
#                                            row_labels = consist_diff_genes, 
#                                            column_title_rot = 90, 
#                                            name = 'log2FC', use_raster = TRUE,
#                                            row_title_rot = 0)
# ComplexHeatmap::draw(consist_diff_hm, padding = unit(c(2, 2, 32, 2), "mm"), row_title = "Marker Genes")

# write consist diff file
consist_diff %>% left_join(consist_diff %>% group_by(Symbol) %>% summarise(Count = n())  %>% left_join(consist_diff %>%  group_by(Symbol) %>% slice_max(., n=1, order_by = `mean log2FC`), by = 'Symbol') %>% select(Symbol,Base, `CT Diff Count` = Count), by = c('Symbol','Base')) %>% mutate(ProposedMarkerGene = case_when(`CT Diff Count` <= 3 ~ 'Yes', TRUE ~ 'No')) %>% arrange(Base, -`mean log2FC`) %>% write_csv('supplemental_files/consistently_differentially_expressed_genes.csv.gz')

toc()


################


###############

tic()

pairwise_neurogenic_test_ungrouped <-  scEiaD_2020_v01 %>% 
  tbl("diff_testing") %>% 
  filter(Against  == "RPC",   
         Group == 'CellType (Predict)', 
         Base == "Neurogenic Cell") %>% 
  #group_by(Base, Against, Gene) %>% 
  #summarise(`mean log2FC` = mean(log2FoldChange), `mean padj` = mean(padj), `sig count` = sum(pvalue < 1e-2), `na count` = sum(is.na(log2FoldChange))) %>% 
  #arrange(-`mean log2FC`) %>%   
  collect() 


pairwise_neurogenic_test<-  scEiaD_2020_v01 %>% 
  tbl("diff_testing") %>% 
  filter(Against  == "RPC",   
         Group == 'CellType (Predict)', 
         Base == "Neurogenic Cell") %>% 
  group_by(Base, Against, Gene) %>% 
  summarise(`mean log2FC` = mean(log2FoldChange), `mean padj` = mean(padj), `sig count` = sum(pvalue < 0.05), `na count` = sum(is.na(log2FoldChange)), baseMean = mean(baseMean), lfcSE = mean(lfcSE)) %>% 
  arrange(-`mean log2FC`) %>%   
  collect() %>% 
  filter(!is.na(`mean log2FC`))


# mutate(GENEID = str_extract(Gene, 'ENS\\w+')) %>% 
# left_join(gene_tab, by = 'GENEID') %>% 
# relocate(SYMBOL, GENEID) %>% 
# dplyr::rename(Symbol = SYMBOL, ID = GENEID) %>% 
# arrange(-`mean log2FC`)

down <- pairwise_neurogenic_test %>% filter(`sig count` > 1) %>% arrange(`mean padj`) %>% filter(`mean log2FC` < 0) %>% head(5) %>% pull(Gene)
up <-  pairwise_neurogenic_test %>% filter(`sig count` > 1) %>% arrange(`mean padj`) %>% filter(`mean log2FC` > 0) %>% head(5) %>% pull(Gene)
source('src/make_exp_plot.R')

input <- list()
input$exp_plot_groups <- c('study_accession')
input$exp_plot_facet <- c('CellType_predict')
input$exp_plot_genes <- down
input$exp_filter_cat <- 'CellType_predict'
#input$exp_filter_on <- c('AC/HC Precursor', 'Amacrine Cell', 'Astrocyte', 'Bipolar Cell', 'Cone', 'Early RPC',' Horizontal Cell','Late RPC','Microglia','Muller Glia', 'Neurogenic Cell', 'Photoreceptor Precursor','Retinal Ganglion Cell','Rod','Rod Bipolar Cell','RPC','RPE' )
input$exp_filter_on <- c("Neurogenic Cell", "RPC" )
input$exp_plot_ylab <- 'Mean log2(Counts + 1)'
#input$exp_plot_ylab <- '% of Cells Detected'
input$exp_plot_col_num <- 16
input$exp_filter_min_cell_number <- 50
input$flip_facet <- TRUE

exp_plot_giga<- make_exp_plot(input, scEiaD_2020_v01, meta_filter) 

input$exp_plot_genes <- up
exp_plot_iovs <- make_exp_plot(input, scEiaD_2020_v01, meta_filter)
a_legend <- get_legend(exp_plot_giga)
# plot_grid(plot_grid(exp_plot_giga + theme(legend.position="none"),
#                     exp_plot_iovs + theme(legend.position="none") + ylab(''), labels = c('a','b')),
#           a_legend, ncol = 1, rel_heights = c(1,0.2))
# toc()


#########################


#########################


tic()
# flip_flop diff genes
down_tab <- down_tab_big %>% 
  mutate(gene_ct_combo = paste(Symbol, Base, sep = '_')) %>% 
  filter(Base %in% ws_oct, 
         Against == 'All')

org_diff_diff_genes <- diff_tab_big %>% 
  mutate(gene_ct_combo = paste(Symbol, Base, sep = '_')) %>% 
  filter(gene_ct_combo %in% down_tab$gene_ct_combo, 
         Against == 'All') %>% 
  arrange(padj) %>% mutate(x = paste0(Symbol, ' (', ID, ')')) 

flip_flop_genes <- scEiaD_2020_v01 %>% tbl('diff_testing_genes') %>% collect() %>% filter(grepl(org_diff_diff_genes %>% pull(ID) %>% paste0(., collapse = '|'), Gene)) %>% pull(Gene)
flip_flop_long <- scEiaD_2020_v01 %>%
  tbl("diff_testing") %>%
  filter(Gene %in% flip_flop_genes) %>%
  collect() %>%
  mutate(Symbol = str_extract(Gene, '^\\w+ ') %>% gsub(' $', '', .)) %>% 
  mutate(gene_ct_combo = paste(Symbol, Base, sep = '_')) %>% 
  filter(gene_ct_combo %in% org_diff_diff_genes$gene_ct_combo,
         Group == 'CellType (Predict)',
         Against == 'All') %>%
  select(Gene, Organism, log2FoldChange, Base)

flip_flop_wide <- flip_flop_long %>% pivot_wider(values_from = log2FoldChange, names_from = Organism)
flip_flop_organism <- colnames(flip_flop_wide)[c(3,4,5)]
flip_flop_wide <- flip_flop_wide %>% data.frame()
row.names(flip_flop_wide) <- paste0(flip_flop_wide$Gene,'_', flip_flop_wide$Base)

flip_flop_wide <- flip_flop_wide[,-c(1,2)]


flip_flop_genes <- row.names(flip_flop_wide) %>% gsub(' \\(.*','',.)

flip_flop_base <- row.names(flip_flop_wide) %>% gsub('.*_','',.) 


# hm_flipflop <- ComplexHeatmap::Heatmap(flip_flop_wide, column_labels = flip_flop_organism, 
#                                        row_split = flip_flop_base, show_row_dend = FALSE, 
#                                        row_labels = flip_flop_genes, 
#                                        column_title_rot = 90, 
#                                        name = 'log2FC', use_raster = TRUE,
#                                        row_title_rot = 0)
# hm_flipflop
toc()

save.image(file = 'data/diff_data_pull.Rdata')
```
