## Supplementary material for "PLAE web app enables powerful searching and multiple visualizations across one million unified single-cell ocular transcriptomes": R scripts: ensdb.R

```
# fix  occasional NA
library(org.Hs.eg.db)
library(EnsDb.Hsapiens.v86)
ensembl_id <- keys(EnsDb.Hsapiens.v86, keytype = "GENEID")
ens_sym_table <- ensembldb::select(EnsDb.Hsapiens.v86, keys= ensembl_id, keytype = "GENEID", columns = c("SYMBOL","GENEID"))
write_tsv(ens_sym_table, file = 'data/hs.ensdb.v86.symbol.geneid.tsv.gz')


library(org.Mm.eg.db)
library(EnsDb.Mmusculus.v79)
ensembl_id <- keys(EnsDb.Mmusculus.v79, keytype = "GENEID")
ens_sym_table <- ensembldb::select(EnsDb.Mmusculus.v79, keys= ensembl_id, keytype = "GENEID", columns = c("SYMBOL","GENEID"))
write_tsv(ens_sym_table, file = 'data/mm.ensdb.v79.symbol.geneid.tsv.gz')


hs_mm <- bind_rows(hs_v86_tab, mm_v79_tab)

diff_tab_big <- scEiaD_2020_v01 %>% tbl('diff_testing') %>% 
  filter(Group == 'CellType (Predict)', padj < 1e-5, (log2FoldChange) > 2) %>% 
  collect() %>% 
  separate(Gene, into = c('Symbol', 'ID'), sep = ' ') %>% 
  mutate(ID = gsub('\\(|\\)', '', ID))

diff_tab_big2 <- diff_tab_big %>% 
  filter(Symbol == '-' | Symbol == 'NA')

diff_tab_big2 <- 
  diff_tab_big2 %>% 
  left_join(hs_mm, by = c('ID' = 'GENEID')) %>% 
  select(-Symbol) %>% 
  rename(Symbol = SYMBOL) %>% 
  relocate(Symbol)

diff_tab_big <- bind_rows(
  diff_tab_big %>% filter(!Symbol %in% c('-', 'NA')),
  diff_tab_big2
)

write_tsv(diff_tab_big, file = 'data/diff_tab_big.tsv.gz')


# down tab

down_tab_big <- scEiaD_2020_v01 %>% tbl('diff_testing') %>% 
  filter(Group == 'CellType (Predict)', padj < 1e-5, (log2FoldChange) < -2) %>% 
  collect() %>% 
  separate(Gene, into = c('Symbol', 'ID'), sep = ' ') %>% 
  mutate(ID = gsub('\\(|\\)', '', ID))

down_tab_big2 <- down_tab_big %>% 
  filter(Symbol == '-' | Symbol == 'NA')

down_tab_big2 <- 
  down_tab_big2 %>% 
  left_join(hs_mm, by = c('ID' = 'GENEID')) %>% 
  select(-Symbol) %>% 
  rename(Symbol = SYMBOL) %>% 
  relocate(Symbol)

down_tab_big <- bind_rows(
  down_tab_big %>% filter(!Symbol %in% c('-', 'NA')),
  diff_tab_big2
)

write_tsv(down_tab_big, file = 'data/down_tab_big.tsv.gz')
```
