## Supplementary material for "PLAE web app enables powerful searching and multiple visualizations across one million unified single-cell ocular transcriptomes": R scripts: make_exp_plot.R

```
make_exp_plot <- function(input, db, meta_filter){
  # TESTING VALUES
  # db <- scEiaD_2020_v01
  # input <- list()
  # input$exp_filter_cat <- 'CellType_predict'
  # input$exp_filter_on <- c('Rods' ,'Cones')
  # input$exp_plot_facet <- c('CellType_predict')
  # input$exp_plot_genes <- c('PAX6 (ENSG00000007372)', 'CRX (ENSG00000105392)', 'NRL (ENSG00000129535)', 'RHO (ENSG00000163914)')
  # input$exp_plot_groups <- c('Stage')
  # input$exp_plot_ylab <- 'Mean log2(Counts + 1)'
  # input$exp_plot_col_num <- 10
  
  gene <- input$exp_plot_genes
  
  grouping_features <- input$exp_plot_groups
  
  if (!'figure_bloop' %in% names(input)){
    input$figure_bloop <- FALSE
  } 
  
  meta_filter_EXP <- meta_filter
  if (!is.null(input$exp_filter_cat)){
    # box_data <- db %>% tbl('grouped_stats') %>%
    #   filter(Gene %in% gene) %>%
    #   collect() %>%
    #   filter(!!as.symbol(input$exp_filter_cat) %in% input$exp_filter_on)
    #
    
    if (class(input$exp_filter_on) == 'character'){
      box_data <- db %>% tbl('grouped_stats') %>%
        filter(Gene %in% gene) %>%
        collect() %>%
        filter_at(vars(all_of(input$exp_filter_cat)), all_vars(. %in% input$exp_filter_on))
      meta_filter_EXP <- meta_filter %>%
        filter_at(vars(all_of(input$exp_filter_cat)), all_vars(. %in% input$exp_filter_on))
      #filter(!!as.symbol(input$exp_filter_cat) %in% input$exp_filter_on)
    } else {
      box_data <- db %>% tbl('grouped_stats') %>%
        filter(Gene %in% gene) %>%
        collect() %>%
        #filter(!!as.symbol(input$exp_filter_cat) %in% input$exp_filter_on) %>%
        filter(!!as.symbol(input$exp_filter_cat) >= input$exp_filter_on[1],
               !!as.symbol(input$exp_filter_cat) <= input$exp_filter_on[2])
      meta_filter_EXP <- meta_filter %>%
        filter(!!as.symbol(input$exp_filter_cat) >= input$exp_filter_on[1],
               !!as.symbol(input$exp_filter_cat) <= input$exp_filter_on[2])
    }
    box_data <- box_data %>% 
      filter(!is.na(!!as.symbol(input$exp_filter_cat))) 
  } else {
    box_data <- db %>% tbl('grouped_stats') %>%
      filter(Gene %in% gene) %>%
      collect()
  }
  # validate(
  #   need(input$exp_plot_groups != '', "Please select at least one grouping feature")
  # )
  
  
  
  #cat(input)
  box_data_temp <- list()
  box_data_NA <- list()
  for (i in gene){
    #print(i)
    box_data_temp[[i]] <- box_data %>%
      filter(Gene == i) %>% 
      filter(!Platform %in% c('SCRBSeq')) %>%
      # mutate(CellType_predict = case_when(CellType_predict == 'RPC' ~ 'RPCs',
      #                                     CellType_predict == 'Mesenchymal/RPE/Endothelial' ~ 'Endothelial',
      #                                     TRUE ~ CellType_predict)) %>%
      mutate(Stage = factor(Stage, levels = c('Early Dev.', 'Late Dev.', 'Maturing', 'Mature'))) %>%
      #filter(!is.na(!!as.symbol(grouping_features))) %>%
      group_by_at(vars(one_of(c('Gene', input$exp_plot_facet, grouping_features)))) %>%
      summarise(counts = sum(counts * cell_exp_ct) / sum(cell_exp_ct),
                cell_exp_ct = sum(cell_exp_ct, na.rm = TRUE)) %>%
      full_join(., meta_filter_EXP %>%
                  group_by_at(vars(one_of(input$exp_plot_facet, grouping_features))) %>%
                  summarise(Count = n())) %>%
      mutate(cell_exp_ct = ifelse(is.na(cell_exp_ct), 0, cell_exp_ct)) %>%
      mutate(`%` = round((cell_exp_ct / Count) * 100, 2),
             Expression = round(counts * (`%` / 100), 2)) %>%
      select_at(vars(one_of(c('Gene', grouping_features, 'cell_exp_ct', 'Count', '%', 'Expression')))) %>%
      arrange(-Expression) %>%
      rename(`Cell # Detected` = cell_exp_ct,
             `Total Cells` = Count,
             `Mean log2(Counts + 1)` = Expression,
             `% of Cells Detected` = `%`) %>%
      
      mutate(`Mean log2(Counts + 1)` = case_when(is.na(`Mean log2(Counts + 1)`) ~ 0, TRUE ~ `Mean log2(Counts + 1)`)) %>% 
      filter(`Total Cells` > as.integer(input$exp_filter_min_cell_number))
    # expand missing genes (nothing detected) to all genes used
    box_data_NA[[i]] <-  box_data_temp[[i]] %>% filter(is.na(Gene))
    if (nrow(box_data_NA[[i]] > 0)){
      box_data_temp[[i]] <- bind_rows(box_data_temp[[i]] %>% filter(!is.na(Gene)),
                                      box_data_NA[[i]] %>% mutate(Gene = i))
    }
  }
  box_data <- box_data_temp %>% bind_rows()
  
  
  box_data$Group <- box_data[,c(2:(length(grouping_features)+1))] %>% tidyr::unite(x, sep = ' ') %>% pull(1)
  
  if (input$flip_facet){
    box_data  %>% 
      mutate(Gene = str_extract(Gene, '^\\w+ ') %>% gsub(" $", '', .)) %>%
      ggplot(aes(x=!!as.symbol(input$exp_plot_facet), y = !!as.symbol(input$exp_plot_ylab), color = !!as.symbol(grouping_features))) +
      geom_boxplot(color = 'black', outlier.shape = NA) +
      ggbeeswarm::geom_quasirandom(size = 4, groupOnX = TRUE) +
      cowplot::theme_cowplot() +
      theme(axis.text.x = element_text(angle = 90, vjust = 0.5, hjust=1)) +
      scale_radius(range=c(2, 6)) +
      scale_colour_manual(values = rep(c(pals::alphabet() %>% unname()), 20)) +
      theme(legend.position="bottom") +
      facet_wrap(ncol = as.numeric(input$exp_plot_col_num), scales = 'free', 'Gene') 
  } else if (input$figure){
    box_data  %>% 
      mutate(Gene = str_extract(Gene, '^\\w+ ') %>% gsub(" $", '', .)) %>%
      ggplot(aes(x=!!as.symbol(input$exp_plot_facet), y = !!as.symbol(input$exp_plot_ylab), color = !!as.symbol(grouping_features), group = CellType_predict, shape = organism)) +
      geom_boxplot(color = 'black', outlier.shape = NA) +
      ggbeeswarm::geom_quasirandom(size = 4, groupOnX = TRUE) +
      cowplot::theme_cowplot() +
      theme(axis.text.x = element_text(angle = 90, vjust = 0.5, hjust=1)) +
      scale_radius(range=c(2, 6)) +
      scale_colour_manual(values = rep(c(pals::alphabet() %>% unname()), 20)) +
      theme(legend.position="bottom") +
      facet_wrap(ncol = as.numeric(input$exp_plot_col_num), scales = 'free', 'Gene') 
  }  else if (input$figure_fovea){
    box_data  %>% 
      filter(!is.na(retina_region)) %>% 
      mutate(Gene = str_extract(Gene, '^\\w+ ') %>% gsub(" $", '', .)) %>%
      ggplot(aes(x=!!as.symbol(input$exp_plot_facet), y = !!as.symbol(input$exp_plot_ylab), color = !!as.symbol(grouping_features), group = CellType_predict)) +
      geom_boxplot(color = 'black', outlier.shape = NA) +
      ggbeeswarm::geom_quasirandom(size = 4, groupOnX = TRUE) +
      cowplot::theme_cowplot() +
      theme(axis.text.x = element_text(angle = 90, vjust = 0.5, hjust=1)) +
      scale_radius(range=c(2, 6)) +
      scale_colour_manual(values = rep(c(pals::alphabet() %>% unname()), 20)) +
      theme(legend.position="bottom") +
      facet_wrap(ncol = as.numeric(input$exp_plot_col_num), ~Gene + retina_region) 
  }    else if (input$figure_bloop){
    box_data  %>% 
      mutate(Gene = str_extract(Gene, '^\\w+ ') %>% gsub(" $", '', .)) %>%
      ungroup() %>% 
      ggplot(aes(x=!!as.symbol(input$exp_plot_facet), y = !!as.symbol(input$exp_plot_ylab), color = !!as.symbol(grouping_features), group = CellType_predict)) +
      geom_boxplot(color = 'black', outlier.shape = NA) +
      ggbeeswarm::geom_quasirandom(size = 1, groupOnX = TRUE) +
      cowplot::theme_cowplot() +
      theme(axis.text.x = element_text(angle = 90, vjust = 0.5, hjust=1)) +
      scale_radius(range=c(2, 6)) +
      scale_colour_manual(values = rep(c(pals::alphabet() %>% unname()), 20)) +
      theme(legend.position="bottom") +
      facet_wrap(ncol = 1, ~Gene, scales = 'free_y') 
  }
  else {
    box_data %>%
      mutate(Gene = str_extract(Gene, '^\\w+ ') %>% gsub(" $", '', .)) %>%
      ggplot(aes(x=Gene, y = !!as.symbol(input$exp_plot_ylab), color = !!as.symbol(grouping_features))) +
      geom_boxplot(color = 'black', outlier.shape = NA) +
      ggbeeswarm::geom_quasirandom(size = 4, groupOnX = TRUE) +
      cowplot::theme_cowplot() +
      theme(axis.text.x = element_text(angle = 90, vjust = 0.5, hjust=1)) +
      scale_radius(range=c(2, 6)) +
      scale_colour_manual(values = rep(c(pals::alphabet() %>% unname()), 20)) +
      theme(legend.position="bottom") +
      facet_wrap(ncol = as.numeric(input$exp_plot_col_num), scales = 'free_x', vars(!!as.symbol(input$exp_plot_facet)))
  }
}
```
