## Supplementary material for "PLAE web app enables powerful searching and multiple visualizations across one million unified single-cell ocular transcriptomes": R scripts: new_fig1_test.R

```
study_count_by_tissue <- meta_filter %>%
  mutate(Organ = case_when(Organ == 'Eye' ~ 'Eye',
                           TRUE ~ 'Body'),
         Tissue = gsub('\\.|_', ' ', Tissue),
         Tissue = gsub("RPE$","RPE-Choroid", Tissue),
         Tissue = gsub(" and Aorta","", Tissue)) %>% 
  #filter(Organ == 'Eye') %>% 
  select(Organ, Tissue,organism, study_accession) %>%
  unique() %>%
  group_by(Organ, Tissue, organism) %>% 
  summarise(Count = n()) %>%
  ungroup() %>% 
  ggplot(aes(x=Tissue,y=Count, fill = organism, label = Count, group = organism)) +
  ylab('Number of Studies') +
  geom_bar(stat='identity', position='stack') +
  shadowtext::geom_shadowtext(position = position_stack(vjust = 0.5)) +
  cowplot::theme_cowplot() +
  coord_flip() +
  theme(legend.position = "none") +
  scale_y_continuous(expand = c(0, 0))  + xlab('') +
  ggforce::facet_col(~Organ, scales = 'free_y', space = 'free')

# retina cell types
retina_ct <- meta_filter %>% filter(Organ == 'Eye') %>% 
  pull(CellType) %>% unique()

cell_counts <- meta_filter %>% filter(Organ == 'Eye', CellType_predict %in% retina_ct) %>% 
  mutate(CellType_predict = case_when(is.na(CellType_predict) ~ 'Unlabelled', TRUE ~ CellType_predict)) %>% 
  group_by(CellType_predict, organism) %>%
  summarise(Count = n()) %>% 
  mutate(sCount = log2(Count)) %>% 
  ggplot(aes(x=CellType_predict,y=sCount, label = Count, fill = organism, group = organism)) +
  ylab('log2(Number of Cells)') +
  geom_bar(stat='identity', position='stack') +
  shadowtext::geom_shadowtext(position = position_stack(vjust = 0.5)) + 
  cowplot::theme_cowplot() +
  coord_flip() + 
  scale_y_continuous(expand = c(0, 0)) + xlab('')

study_counts_by_cell <- meta_filter %>% filter(Organ == 'Eye', CellType_predict %in% retina_ct) %>% 
  mutate(CellType_predict = case_when(is.na(CellType_predict) ~ 'Unlabelled', TRUE ~ CellType_predict)) %>% 
  group_by(organism, CellType_predict, study_accession) %>%
  summarise(Count = n()) %>% 
  filter(Count > 10) %>% 
  summarise(Count = n()) %>% 
  mutate(sCount = log2(Count)) %>% 
  ggplot(aes(x=CellType_predict,y=Count, label = Count, fill = organism, group = organism)) +
  ylab('Number of Studies') +
  geom_bar(stat='identity', position='stack') +
  shadowtext::geom_shadowtext(position = position_stack(vjust = 0.5)) + 
  cowplot::theme_cowplot() +
  coord_flip() + 
  scale_y_continuous(expand = c(0, 0)) + xlab('')

timings <- rbind(
  c(1,7,1),
  c(9,13,8),
  c(3,NA,2),
  c(2,NA,NA),
  c(2,1,NA),
  c(2,NA,NA),
  c(2,1,NA),
  c(4,41,NA)
) %>% data.frame()

row.names(timings) <- c('Initial Page',
                        'Cell Type Colored UMAP',
                        'Abca4 Expression UMAP',
                        'Boxplot of Abca4 Expression By CellType Across All Studies',
                        'Boxplot of Abca4 Expression By CellType Across One Study',
                        'Dotplot of Abca4, Pax6, Nrl, and Atoh7',
                        'Heatmap of Abca4, Pax6, Nrl, and Atoh7',
                        'Table of Differentially Expressed Genes between Bipolar and Cone'
                        
)

colnames(timings) <- c('PLAE',"Spectacle",'Tabula Sapiens\n(CELLxGENE)')

timings %>% as_tibble(rownames = 'Test') %>% 
  pivot_longer(-Test) %>% 
  mutate(Test = factor(Test, levels = row.names(timings))) %>% 
  ggplot(aes(x=name,y=value)) + 
  geom_bar(stat= 'identity') + 
  facet_wrap(~Test, scales = 'free', ncol = 2) + 
  xlab('Resource') + 
  ylab('Seconds to Load') + 
  cowplot::theme_cowplot()


timings %>% as_tibble(rownames = 'Test') %>% 
  pivot_longer(-Test) %>% 
  mutate(Test = factor(Test, levels = row.names(timings))) %>% 
  ggplot(aes(x=name,y=value)) + 
  geom_bar(stat= 'identity') + 
  facet_grid(~Test) #, scales = 'free', ncol = 2) + 
xlab('Resource') + 
  ylab('Seconds to Load') + 
  cowplot::theme_cowplot()
```
