## Supplementary material for "PLAE web app enables powerful searching and multiple visualizations across one million unified single-cell ocular transcriptomes": R scripts: pubmed_query.R

```
# return PMID for query
library(easyPubMed)
library(tidyverse)
#load('~/data/massive_integrated_eye_scRNA/top_markers.Rdata')
library(pool)
library(RSQLite)

#data_dir_vPLAE <- '/Volumes/McGaughey_S/data/scEiaD_2022_02/'
#scEiaD_2020_v01 <- pool::dbPool(RSQLite::SQLite(), dbname =  glue('{data_dir_vPLAE}/MOARTABLES__anthology_limmaFALSE___4000-counts-universe-study_accession-scANVIprojection-15-5-0.1-50-20__pointRelease01.sqlite'))

pubmed_counter <- function(query, sleep_time = 0.2, api_key = '8d33dae565ca3a523495197da36cc085d308'){
  Sys.sleep(sleep_time)
  pub_query <- query
  entrez_id <- get_pubmed_ids(pub_query, api_key = api_key)
  if (entrez_id$Count == 0){
    # run one more time, with a 0.5 second delay
    Sys.sleep(1)
    print("RERUN")
    entrez_id <- get_pubmed_ids(pub_query, api_key = api_key)
  }
  if (entrez_id$Count == 0){
    # then give up and return NA
    print("NA")
    out <- NA
  } else {
    abstracts_txt <- fetch_pubmed_data(entrez_id, format = "abstract") 
    out <- grep('PMID', abstracts_txt, value = TRUE) %>% str_extract(., '\\d\\d\\d+') %>% unique()}
  out
}

#ctr <- meta_filter %>% filter(Tissue == 'Retina') %>% pull(CellType)

queries <- consist_diff %>% 
  #filter(Symbol %in% consist_diff_genes) %>% 
  #filter(Base %in% ctr) %>% 
  mutate(Base2 = gsub('Cell','',Base),
         Base2 = case_when(Base2 == 'AC/HC Precursor' ~ 'Amacrine Horizontal',
                          grepl('RPC', Base2) ~ 'Progenitor',
                          TRUE ~ Base2)) %>% 
  mutate(Symbol2 = case_when(Symbol == 'TF' ~ 'Transferrin',
                            Symbol == 'CP' ~ 'ceruloplasmin',
                            Symbol == 'F3' ~ 'Coagulation Factor III',
                            Symbol == 'HR' ~ 'hairless',
                            TRUE ~ Symbol)) %>% 
  mutate(
    pm_query0 = paste0(Symbol2),
    pm_query1 = paste0(Symbol2, ' AND ', Base2),
    pm_query2 = paste0(Symbol2, ' AND Retina'))

# heatmap_fig_genes <- list()
# for (i in consist_diff_genes){
#   print(i)
#   heatmap_fig_genes[[i]] <- pubmed_counter(i)
# }

pmid0 <- list()
for (i in unique(queries$pm_query0)){
  print(i)
  pmid0[[i]] <- pubmed_counter(i)
}

pmid1 <- list()
for (i in unique(queries$pm_query1)){
  print(i)
  #Sys.sleep(1)
  pmid1[[i]] <- pubmed_counter(i)
}

pmid2 <- list()
for (i in unique(queries$pm_query2)){
  print(i)
  #Sys.sleep(1)
  pmid2[[i]] <- pubmed_counter(i)
}


set.seed(1423)
all_genes <- scEiaD_2020_v01 %>% tbl('Genes') %>% pull(Gene) %>% gsub(' \\(.*','', .)
all_diff_symbol <- diff_tab %>% filter(padj < 0.01) %>% pull(Symbol) %>% unique()


rand_not_marker0 <- all_genes[!all_genes %in% queries$Symbol] 
rand_not_marker1 <- all_genes[!all_genes %in% all_diff_symbol] 

rand_not_marker0_sample <- rand_not_marker0[sample(1:length(rand_not_marker0), 1000)]
rand_not_marker1_sample <- rand_not_marker1[sample(1:length(rand_not_marker1), 1000)]
rand_all <- all_genes[sample(1:length(all_genes), 1000)]

pm_query3 <- paste0(rand_not_marker1_sample, ' AND Retina')
pmid3 <- list()
for (i in pm_query3){
  print(i)
  #Sys.sleep(1)
  pmid3[[i]] <- pubmed_counter(i)
}

pm_query4 <- rand_not_marker1_sample
pmid4 <- list()
for (i in pm_query4){
  print(i)
  #Sys.sleep(1)
  pmid4[[i]] <- pubmed_counter(i)
}

pm_query5 <- rand_all
pmid5 <- list()
for (i in pm_query5){
  print(i)
  #Sys.sleep(1)
  pmid5[[i]] <- pubmed_counter(i)
}

pm_query6 <- paste0(rand_all, ' AND Retina')
pmid6 <- list()
for (i in pm_query6){
  print(i)
  #Sys.sleep(1)
  pmid6[[i]] <- pubmed_counter(i)
}
# pmid2 <- list()
# for (i in x$pm_query2){
#   print(i)
#   #Sys.sleep(1)
#   pmid2[[i]] <- pubmed_counter(i)
# }


queries <- queries %>% left_join(., pmid0 %>% map(function(x) sum(!is.na(x))) %>% unlist() %>% enframe() %>% rename(query0=value), by = c("Symbol" = "name")) 
queries <- queries %>% left_join(., pmid1 %>% map(function(x) sum(!is.na(x))) %>% unlist() %>% enframe() %>% rename(query1=value), by = c("pm_query1" = "name")) 
queries <- queries %>% left_join(., pmid2 %>% map(function(x) sum(!is.na(x))) %>% unlist() %>% enframe() %>% rename(query2=value), by = c("pm_query2" = "name"))
save(pmid0, pmid1, pmid2, pmid3, pmid4, pmid5, pmid6, queries, file = 'data/top_marker_pmid.Rdata')
```
